# Endosomolytic Peptides Enable the Cellular Delivery of Peptide Nucleic Acids

**DOI:** 10.1101/2024.06.18.599558

**Authors:** JoLynn B. Giancola, Ronald T. Raines

## Abstract

Precision genetic medicine enlists antisense oligonucleotides (ASOs) to bind to nucleic acid targets important for human disease. Peptide nucleic acids (PNAs) have many desirable attributes as ASOs but lack cellular permeability. Here, we use an assay based on the corrective splicing of an mRNA to assess the ability of synthetic peptides to deliver a functional PNA into a human cell. We find that the endosomolytic peptides L17E and L17ER_4_ are highly efficacious delivery vehicles. Co-treatment of a PNA with low micromolar L17E or L17ER_4_ enables robust corrective splicing in nearly all treated cells. Peptide–PNA conjugates are even more effective. These results enhance the utility of PNAs as research tools and potential therapeutic agents.

---

Oligonucleotides have garnered enormous attention for their therapeutic potential. Eighteen nucleic acid-based drugs are on the market, and dozens more are in clinical development.^1^ Because oligonucleotide-target engagement is governed by Watson–Crick–Franklin base pairing, therapeutic oligonucleotides can easily be designed to have high specificity for their target, obviating the need for the iterative screening required to develop small-molecule therapies.^2^

Antisense oligonucleotides (ASOs) are a class of oligonucleotide that leverages mRNA binding as a means to control protein levels in a cell.^3,4^ Upon binding to an mRNA target in the cytosol, an ASO can induce target degradation. Alternatively, binding to an mRNA target in the nucleus can direct splicing that enables (or disables) its translation.

Peptide nucleic acids (PNAs) are heralded ASOs in which the ribose phosphate backbone of an oligonucleotide is replaced with one that is peptidic in nature.^5^ PNAs have exquisite specificity for their complementary nucleic acid and bind with higher affinity than do their DNA counterparts because their neutral backbone obviates the Coulombic repulsion endemic in double-stranded nucleic acids.^6,7^ PNAs are achiral and readily accessible by solid-phase synthesis methods akin to those for accessing peptides.^8^ Moreover, PNAs are highly stable under physiological conditions and not known to be immunogenic. These attributes have made the application of PNAs in the laboratory and clinic of special interest.^9,10^

To function in a live cell, a PNA must enter the cytosol. PNAs are, however, unable to penetrate the plasma membrane of mammalian cells. Their ability to access the cytosol is notoriously inefficient, even with delivery vehicles.^2,11-26^ Accordingly, the utility of PNAs has been constrained since their invention over thirty years ago.^5^

In 2017, Futaki and coworkers reported on L17E, an endosomolytic peptide derived from M-lycotoxin.^27^ This peptide affords cytosolic access to cargo through endosomal membrane rupture after macropinocytotic uptake.^28^ L17E has been employed primarily as a co-treatment agent in protein delivery experiments.^29^ Only a few reports disclose the use of this peptide as a covalent appendage for the delivery of oligonucleotides.^30,31^ In 2022, the Futaki group reported that a pendant arginine-rich sequence makes L17E even more effective for the cellular delivery of proteinaceous cargo.^32^

Here, we assess the efficiency of PNA delivery into human cells as mediated by three peptides: L17E,^27^ L17ER_4_ (which contains four arginine residues),^32^ and R10 (which is a well-known cell-penetrating peptide).^33^ To assess delivery, we employ a rigorous quantitative assay based on the corrective splicing of an mRNA that is translated into the green fluorescent protein (GFP) (Scheme 1). First, we compare the cellular uptake of a PNA upon co-treatment with each peptide. Then, we explore the delivery efficiency upon conjugation of the peptide to the PNA. We find that endosomolytic peptides can effect the highly efficient delivery of a PNA into human cells.

**Scheme 1.**
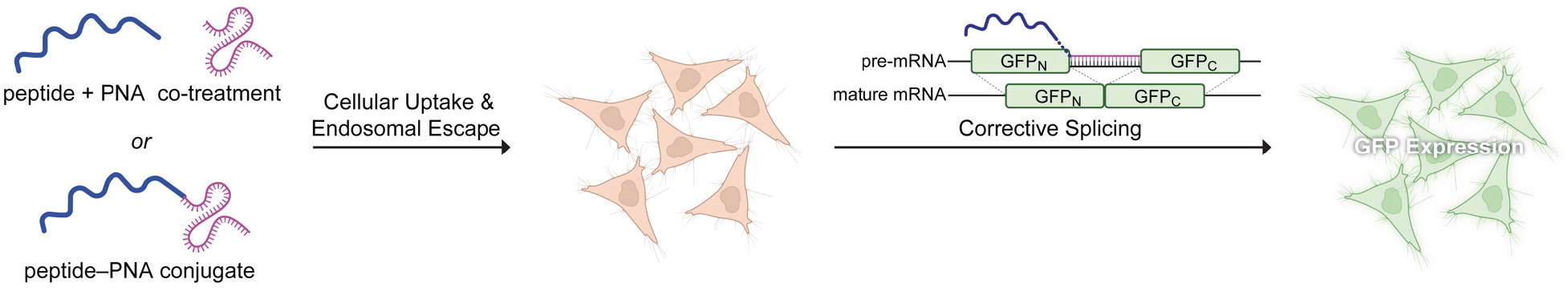
Assay for the Delivery of a Functional PNA into Live Human Cells.

Our first goal was to demonstrate the cytosolic delivery of a PNA into human cells. For this purpose, we chose to deliver PNA654, which can orchestrate corrective splicing of a GFP pre-RNA in an engineered human cell line (HeLa654), resulting in the production of GFP.^34-37^ This assay enabled us to make comparisons between different treatment conditions because GFP fluorescence reports directly on the delivery of a *functional* PNA (Scheme 1). We used standard solid-phase methods to synthesize the necessary peptides (Scheme 2). In some of these peptides, azido groups enable strain-promoted azide–alkyne cycloaddition (SPAAC) for subsequent conjugation.

**Scheme 2.**
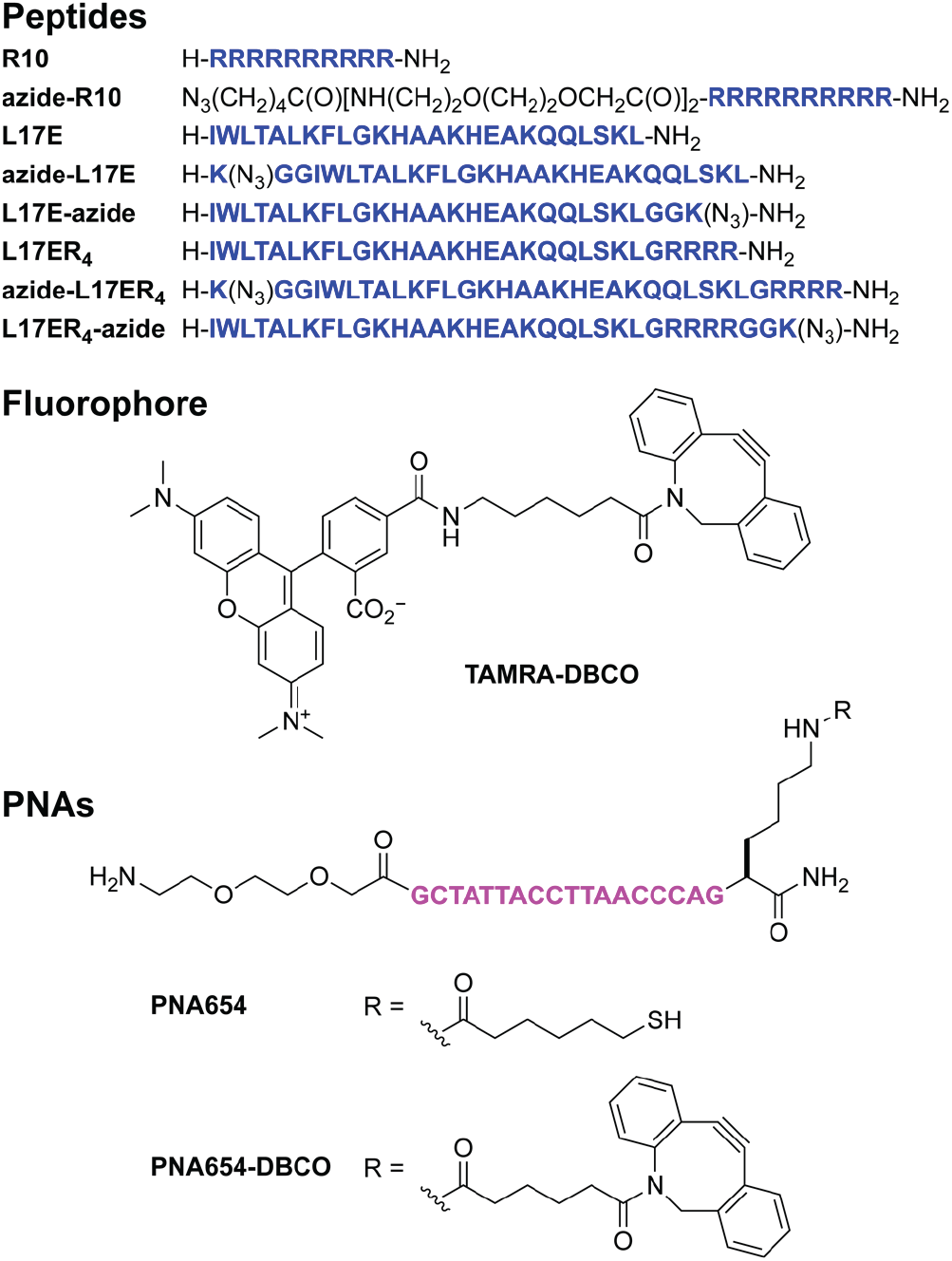
Molecules Used in this Work.

We were aware that cell-penetrating peptides can diminish cell viability.^38^ Accordingly, we treated HeLa654 cells with R10, L17E, or L17ER_4_ and assessed their viability with a tetrazolium-based assay^39^ and their morphology with epifluorescence microscopy. R10 and L17E did not have a detectable effect on viability at all assayed concentrations (Figure 1). In contrast, L17ER_4_ was cytotoxic above 5 µM in the absence of serum and above 20 µM in the presence of serum. These viability data were consistent with our morphological assessments (Figure S8) and provided guidance for subsequent experiments.

**Figure 1.**
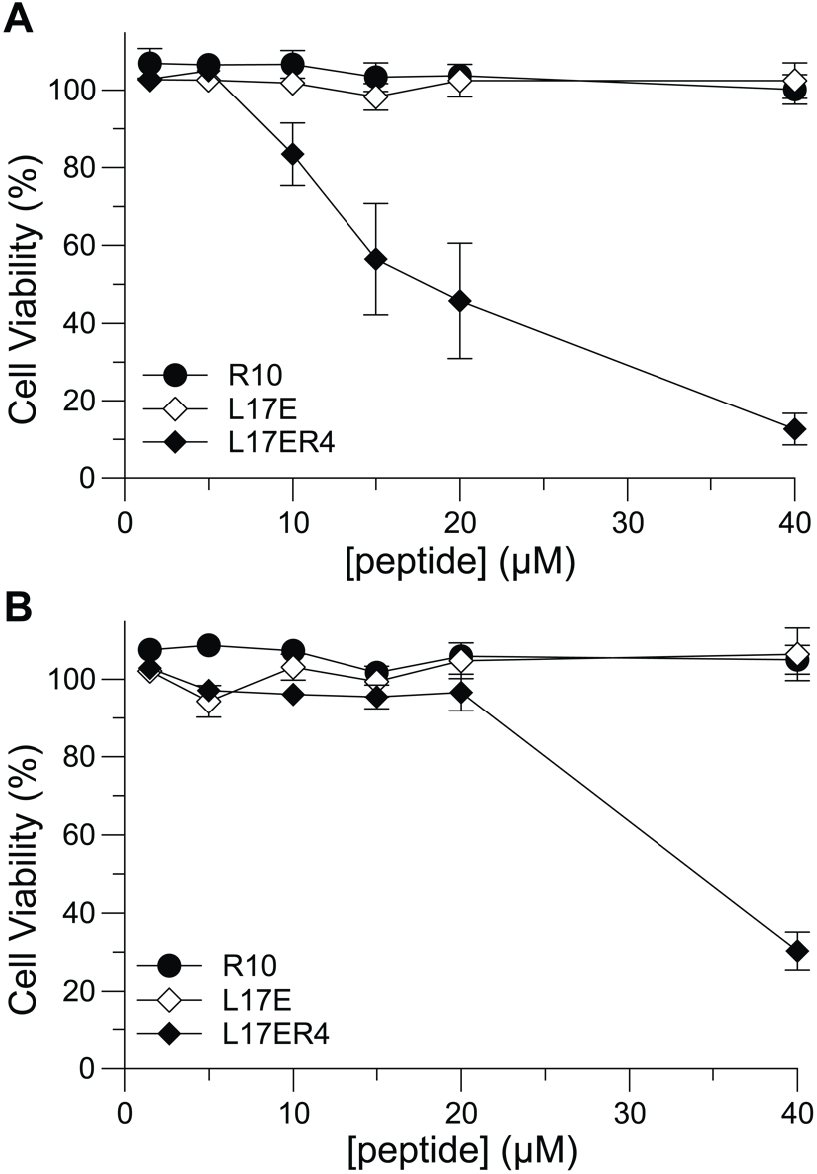
Graphs of HeLa654 cell viability after treatment with R10, L17E, or L17ER_4_. Cells were treated with peptide (1.5 μM– 40 μM) in a humidified incubator at 37 °C for either (A) 5–7 min in serum-free medium or (B) 1 h in complete medium. After treatment, cells were allowed to recover in complete medium until the assay endpoint. Viability was assessed after 16 h. Values are the mean ± SE from two independent experiments, each performed with two technical replicates.

Next, we characterized the cellular uptake of the endosomolytic peptides. To do so, we used SPAAC to link an azido group installed at the N or C terminus with the strained alkynyl group of TAMRA-DBCO. These conjugates also served as proxies for conjugates to larger biomolecules.^32^ Upon incubation with human cells, we found that modification of the endosomolytic peptides at their C terminus led to greater cellular uptake (Figure 2).

**Figure 2.**
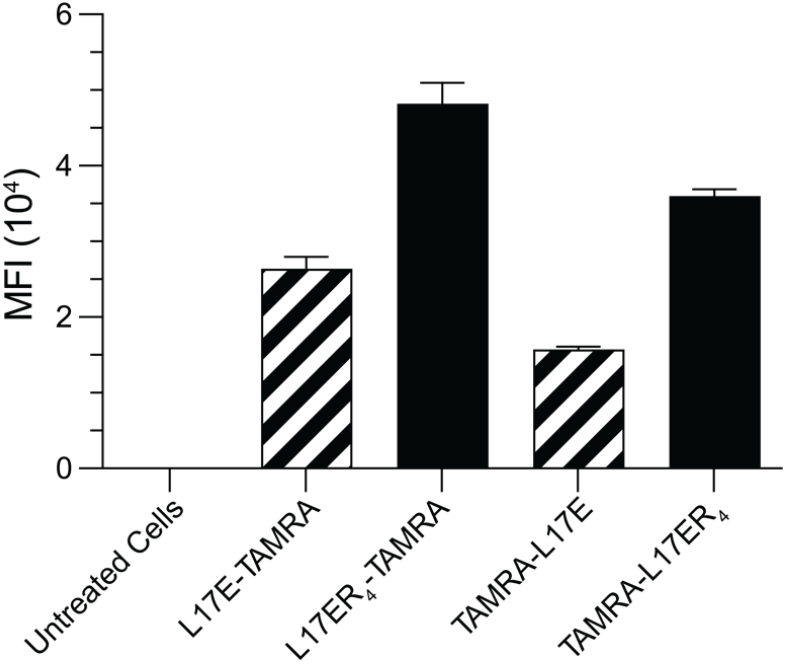
Graphs of the mean fluorescence intensities (MFIs) from TAMRA-labelled L17E and L17ER_4_ within human cells. HeLa654 cells were treated with peptides (5 µM) labeled at their N or C terminus for 5 min in medium without serum, allowed to recover in complete medium for 1 h, and assessed with flow cytometry. For experimental conditions, see Table S2. Values are the mean ± SE from at least two independent experiments, each performed with two technical replicates. Standardized laser intensities were used across all experiments.

Guided by data with TAMRA-labeled peptides, we used SPAAC to prepare the R10–PNA, L17E–PNA, and L17ER_4_– PNA conjugates in which the PNA is installed at the C terminus of the peptide. We then treated HeLa654 cells with these conjugates for 5–7 min in Dulbecco’s modified Eagle’s medium (DMEM) without serum or 1 h in complete medium. These conditions were guided by cellular viability (Figure 1) and literature precedent.^32^ After treatment, the cells were washed with phosphate-buffered saline and permitted to recover for an additional 27 h in complete medium in a humidified incubator at 37 °C prior to epifluorescence imaging.

Our initial experiments on the cellular delivery of a PNA focused on the utility of a co-treatment with R10, L17E, and L17ER_4_. In these experiments, we used R10, L17E, and L17ER_4_ at 40, 40, and 20 µM, respectively, and PNA at 1–30 µM. Without a peptide or with R10, we observed only modest GFP production, indicative of little cellular uptake (Figures 3 and 4). In contrast, we observed substantial fluorescence with 40 µM L17E and, even more so, with 20 µM L17ER_4_. Interestingly, and in contrast to our cytotoxicity data, we observed mild toxicity for cells treated with 20 µM L17ER_4_ and low PNA concentrations (1 and 5 µM), but abrogation of this cytotoxicity with high PNA concentrations (15 and 30 µM). These data suggest that a nearly stoichiometric peptide:PNA ratio is desirable for co-treatments with L17ER_4_ and motivated us to assess a covalent 1:1 conjugate of the peptide and PNA.

**Figure 3.**
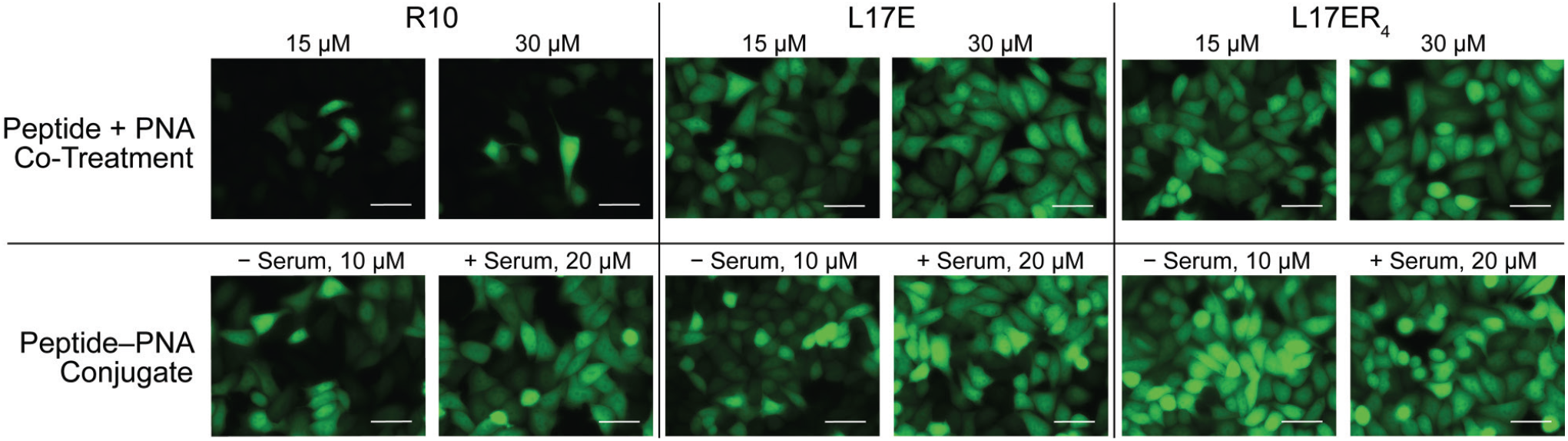
Microscopy images of GFP in human cells produced upon corrective splicing by a PNA delivered with peptide co-treatment or conjugation (Scheme 1). Experimental conditions and additional images are provided in Table S3. Standardized laser intensities were normalized across all treatment conditions. Images are representative of GFP production from at least two independent experiments performed with two technical replicates. Additional Images are shown in Figures S14, S16, S18, S21, S23, and S25. Scale bars: 50 μM.

**Figure 4.**
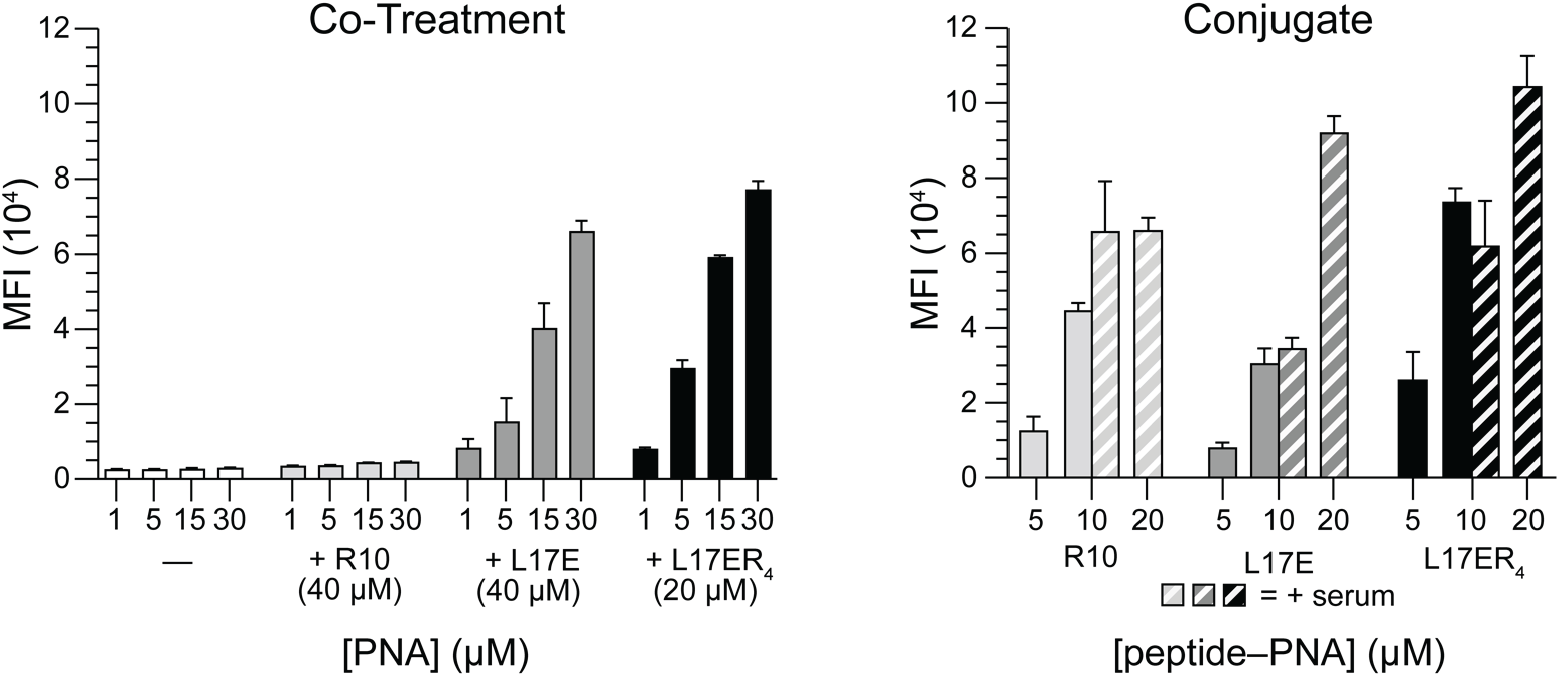
Graphs of MFIs from GFP in human cells produced upon corrective splicing by a PNA delivered with peptide co-treatment or conjugation (Scheme 1). Values are the mean ± SE from at least two independent experiments, each performed with two technical replicates. For experimental conditions, see Table S3. Standardized laser intensities were used across all experiments.

Before testing peptide–PNA conjugates, we probed for an effect of the cell cycle on PNA delivery. To do so, we synchronized cells by using RO-3306, which is a small– molecule CDK1 inhibitor that arrests the cell cycle at the G2/M phase boundary.^40^ We chose that point in the cell cycle because of the known increase in ceramide concentration in membranes there,^41-43^ the role of ceramides in forming nucleation zones that direct endocytosis,^44-50^ and the correlation between improved cargo uptake in dividing cells.^51^ Cells were seeded at equivalent densities, and a subset was synchronized by the addition of RO-3306 (9 µM for 20 h). To ensure that we captured cells after their reentry into the cell cycle but while they were dividing, we administered L17E and the PNA immediately following the withdrawal of RO-3306, using a 2-h treatment window in complete medium. We observed that synchronizing cells did not potentiate GFP production (Figures S27 and S28), suggesting that ceramides are not involved in the uptake mechanism.

Next, we discerned whether covalently linking a peptide to the PNA was beneficial to cellular uptake. Specifically, we treated HeLa654 cells with peptide–PNA conjugates either at 5 and 10 µM for 7 min in DMEM without serum or at 10 and 20 µM for 1 h in complete medium. We observed a marked enhancement in performance by R10 when covalently attached to the PNA (Figure 3). Still, both endosomolytic peptides outperformed R10. Moreover, GFP production appeared to be greater from L17ER_4_–PNA than from L17E–PNA in the absence or presence of serum.

Finally, we used flow cytometry to quantify the cellular GFP induced by a PNA upon co-treatment or conjugation. Consistent with our imaging analysis, co-treatment with R10 affords little GFP (Figures 4 and S18). In contrast, we found a robust dose-dependent response for GFP production upon co-treatment with endosomolytic peptides (Figures 4, S15, and S17). At 1 μM, L17ER_4_ and L17E elicit PNA function to a similar extent, but L17ER_4_ elicits more function than L17E at concentrations ≥5 μM. Notably, we found that at PNA concentrations of 15 and 30 μM, both L17E and L17ER_4_ enable >90% of cells to be transfected successfully with the PNA (Figures S15 and S17). Even at 5 μM L17ER_4_, >85% of cells produce GFP (Figure S17).

A covalent linkage with a peptide enhances the function of the PNA in cells (Figure 4). Under each condition, the GFP levels induced by a conjugate exceeded that obtained with cotreatment. Oftentimes, the cellular delivery of cargo is dampened by the addition of serum because of nonspecific adsorption of the cargo to serum proteins. In marked contrast, the intracellular delivery of PNA conjugates with endosomolytic peptides was potentiated by serum. Moreover, treatment conditions with L17E and L17ER_4_ can enable ∼95% of cells to produce GFP (Figures S22 and S24).

In conclusion, we have found that the endosomolytic peptides L17E and L17ER_4_ are highly efficacious and superior to R10 in effecting the delivery of a functional PNA into a human cell. Unlike with previous delivery strategies, nearly all cells produce GFP after either co-treatment with low micromolar concentrations of L17E or L17ER_4_ or conjugation to these peptides. L17E affords robust cellular uptake without compromising cell viability, with less efficacy than L17ER_4_. The co-treatment strategy is an especially facile means to deliver a PNA into a human cell in vitro, whereas conjugation has the potential for use in vivo. We anticipate that our findings will enhance the utility of PNAs and other ASOs in a wide variety of applications.

## ASSOCIATED CONTENT

### Supporting Information

The Supporting Information is available free of charge on the ACS Publications website at DOI: 10.1021/xxxx.xxxxxxx.

Experimental procedures for the synthesis, characterization, and analysis of peptides; Tables S1–S3; and Figures S1–S28 (PDF)

## AUTHOR INFORMATION

### Author

**JoLynn B. Giancola** – *Department of Chemistry, Massachusetts Institute of Technology, Cambridge, Massachusetts 02139, United States;*

## Notes

The authors declare no competing financial interest.

## ACKNOWLEDGMENTS

The authors are grateful to Dr. Carly K. Schissel and Professor B. L. Pentelute for providing the HeLa654 cell line. They thank the Koch Institute’s Robert A. Swanson (1969) Biotechnology Center for technical support, specifically the High Throughput Sciences core for mycoplasma testing. They are grateful to Dr. Jinyi Yang, Nicola R. F. Knowles, and Dr. Evans C. Wralstad for helpful discussions.

## TOC GRAPHIC

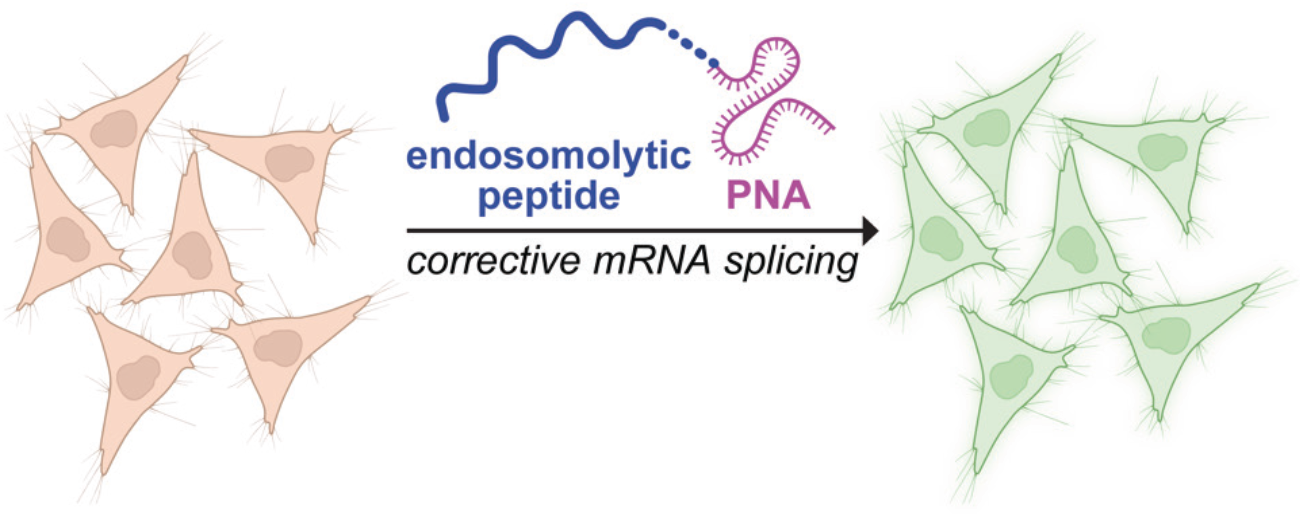

## REFERENCES

(1) Egli, M.; Manoharan, M. Chemistry, structure and function of approved oligonucleotide therapeutics. Nucleic Acids Res. 2023, 51, 2529–2573.

(2) Roberts, T. C.; Langer, R.; Wood, M. J. A. Advances in oligonucleotide drug delivery. Nat. Rev. Drug Discov. 2020, 19, 673–694.

(3) Amantana, A.; Moulton, H. M.; Cate, M. L.; Reddy, M. T.; Whitehead, T.; Hassinger, J. N.; Youngblood, D. S.; Iversen, P. L. Pharmacokinetics, biodistribution, stability and toxicity of a cell-penetrating peptide–morpholino oligomer conjugate. Bioconjugate Chem. 2007, 18, 1325–1331.

(4) Dhuri, K.; Bechtold, C.; Quijano, E.; Pham, H.; Gupta, A.; Vikram, A.; Bahal, R. Antisense oligonucleotides: An emerging area in drug discovery and development. J. Clin. Med. 2020, 9, 2004.

(5) Egholm, M.; Buchardt, O.; Nielsen, P. E.; Berg, R. H. Peptide nucleic acids (PNA). Oligonucleotide analogues with an achiral peptide backbone. J. Am. Chem. Soc. 1992, 114, 1895–1897.

(6) Egholm, M.; Buchardt, O.; Christensen, L.; Behrens, C.; Freier, S. M.; Driver, D. A.; Berg, R. H.; Kim, S. K.; Norden, B.; Nielsen, P. E. PNA hybridizes to complementary oligonucleotides obeying the Watson–Crick hydrogen-bonding rules. Nature 1993, 365, 566–568.

(7) Wittung, P.; Nielsen, P. E.; Buchardt, O.; Egholm, M.; Norden, B. DNA-like double helix formed by peptide nucleic-acid. Nature 1994, 368, 561–563.

(8) Klabenkova, K.; Fokina, A.; Stetsenko, D. Chemistry of Peptide–oligonucleotide conjugates: A review. Molecules 2021, 26, 5420.

(9) Montazersaheb, S.; Hejazi, M. S.; Nozad Charoudeh, H. Potential of peptide nucleic acids in future therapeutic applications. Adv. Pharm. Bull. 2018, 8, 551–563.

(10) Brodyagin, N.; Katkevics, M.; Kotikam, V.; Ryan, C. A.; Rozners, E. Chemical approaches to discover the full potential of peptide nucleic acids in biomedical applications. Beilstein J. Org. Chem. 2021, 17, 1641–1688.

(11) Koppelhus, U.; Nielsen, P. E. Cellular delivery of peptide nucleic acid (PNA). Adv. Drug Deliv. Rev. 2003, 55, 267–280.

(12) Kaihatsu, K.; Huffman, K. E.; Corey, D. R. Intracellular uptake and inhibition of gene expression by PNAs and PNA−peptide conjugates. Biochemistry 2004, 43, 14340–14347.

(13) Turner, J. J.; Ivanova, G. D.; Verbeure, B.; Williams, D.; Arzumanov, A. A.; Abes, S.; Lebleu, B.; Gait, M. J. Cell-penetrating peptide conjugates of peptide nucleic acids (PNA) as inhibitors of HIV-1 Tat-dependent trans-activation in cells. Nucleic Acids Res. 2005, 33, 6837–6849.

(14) Abes, S.; Williams, D.; Prevot, P.; Thierry, A.; Gait, M. J.; Lebleu, B. Endosome trapping limits the efficiency of splicing correction by PNA–oligolysine conjugates. J. Control. Release 2006, 110, 595–604.

(15) Avitabile, C.; Moggio, L.; Malgieri, G.; Capasso, D.; Di Gaetano, S.; Saviano, M.; Pedone, C.; Romanelli, A. γ Sulphate PNA (PNA S): Highly selective DNA binding molecule showing promising antigene activity. PLoS One 2012, 7, e35774.

(16) Spinelli, N.; Defrancq, E.; Morvan, F. Glycoclusters on oligonucleotide and PNA scaffolds: Synthesis and applications. Chem. Soc. Rev. 2013, 42, 4557–4573.

(17) Cordier, C.; Boutimah, F.; Bourdeloux, M.; Dupuy, F.; Met, E.; Alberti, P.; Loll, F.; Chassaing, G.; Burlina, F.; Saison-Behmoaras, T. E. Delivery of antisense peptide nucleic acids to cells by conjugation with small arginine-rich cell-penetrating peptide (R/W)9. PLoS One 2014, 9, e104999.

(18) Avitabile, C.; Accardo, A.; Ringhieri, P.; Morelli, G.; Saviano, M.; Montagner, G.; Fabbri, E.; Gallerani, E.; Gambari, R.; Romanelli, A. Incorporation of naked peptide nucleic acids into liposomes leads to fast and efficient delivery. Bioconjugate Chem. 2015, 26, 1533–1541.

(19) Lehto, T.; Ezzat, K.; Wood, M. J. A.; El Andaloussi, S. Peptides for nucleic acid delivery. Adv. Drug Deliv. Rev. 2016, 106, 172–182.

(20) Gupta, A.; Bahal, R.; Gupta, M.; Glazer, P. M.; Saltzman, W. M. Nanotechnology for delivery of peptide nucleic acids (PNAs). J. Control. Rel. 2016, 240, 302–311.

(21) Saarbach, J.; Sabale, P. M.; Winssinger, N. Peptide nucleic acid (PNA) and its applications in chemical biology, diagnostics, and therapeutics. Curr. Opin. Chem. Biol. 2019, 52, 112–124.

(22) Ghavami, M.; Shiraishi, T.; Nielsen, P. E. Cooperative cellular uptake and activity of octaarginine antisense peptide nucleic acid (PNA) conjugates. Biomolecules 2019, 9, 554.

(23) Shiraishi, T.; Ghavami, M.; Nielsen, P. E. In vitro cellular delivery of peptide nucleic acid (PNA). Methods Mol. Biol. 2020, 2105, 173–185.

(24) Kulkarni, J. A.; Witzigmann, D.; Thomson, S. B.; Chen, S.; Leavitt, B. R.; Cullis, P. R.; van der Meel, R. The current landscape of nucleic acid therapeutics. Nat. Nanotechnol. 2021, 16, 630–643.

(25) Öztürk, Ö.; Lessl, A.-L.; Höhn, M.; Wuttke, S.; Nielsen, P. E.; Wagner, E.; Lächelt, U. Peptide nucleic acid–zirconium coordination nanoparticles. Sci. Rep. 2023, 13, 14222.

(26) Moccia, M.; Pascucci, B.; Saviano, M.; Cerasa, M. T.; Terzidis, M. A.; Chatgilialoglu, C.; Masi, A. Advances in nucleic acid research: Exploring the potential of oligonucleotides for therapeutic applications and biological studies. Int. J. Mol. Sci. 2024, 25, 146.

(27) Akishiba, M.; Takeuchi, T.; Kawaguchi, Y.; Sakamoto, K.; Yu, H. H.; Nakase, I.; Takatani-Nakase, T.; Madani, F.; Gräslund, A.; Futaki, S. Cytosolic antibody delivery by lipid-sensitive endosomolytic peptide. Nat. Chem. 2017, 9, 751–761.

(28) Akishiba, M.; Futaki, S. Inducible membrane permeabilization by attenuated lytic peptides: A new concept for accessing cell interiors through ruffled membranes. Mol. Pharm. 2019, 16, 2540–2548.

(29) Giancola, J. B.; Grimm, J. B.; Jun, J. V.; Petri, Y. D.; Lavis, L. D.; Raines, R. T. Evaluation of the cytosolic uptake of HaloTag using a pH-sensitive dye. ACS Chem. Biol. 2024, 19, 908–915; and references therein.

(30) Becker, B.; Englert, S.; Schneider, H.; Yanakieva, D.; Hofmann, S.; Dombrowsky, C.; Macarrón Palacios, A.; Bitsch, S.; Elter, A.; Meckel, T.; Kugler, B.; Schirmacher, A.; Avrutina, O.; Diederichsen, U.; Kolmar, H. Multivalent dextran hybrids for efficient cytosolic delivery of biomolecular cargoes. J. Peptide Sci. 2021, 27, e3298.

(31) Feng, R.; Ni, R.; Chau, Y. Fusogenic peptide modification to enhance gene delivery by peptide–DNA nano-coassemblies. Biomater. Sci. 2022, 10, 5116–5120.

(32) Shinga, K.; Iwata, T.; Murata, K.; Daitoku, Y.; Michibata, J.; Arafiles, J. V. V.; Sakamoto, K.; Akishiba, M.; Takatani-Nakase, T.; Mizuno, S.; Sugiyama, F.; Imanishi, M.; Futaki, S. L17ER4: A cell-permeable attenuated cationic amphiphilic lytic peptide. Bioorg. Med. Chem. 2022, 61, 116728.

(33) Langel, Ü., CPP, Cell-Penetrating Peptides, 2nd ed. Springer Nature Switzerland: Cham, Switzerland, 2023.

(34) Sazani, P.; Kang, S.-H.; Maier, M. A.; Wei, C.; Dillman, J.; Summerton, J.; Manoharan, M.; Kole, R. Nuclear antisense effects of neutral, anionic and cationic oligonucleotide analogs. Nucleic Acids Res. 2001, 29, 3965–3974.

(35) Wolfe, J. M.; Fadzen, C. M.; Choo, Z.-N.; Holden, R. L.; Yao, M.; Hanson, G. J.; Pentelute, B. L. Machine learning to predict cell-penetrating peptides for antisense delivery. ACS Cent. Sci. 2018, 4, 512–520.

(36) Schissel, C. K.; Mohapatra, S.; Wolfe, J. M.; Fadzen, C. M.; Bellovoda, K.; Wu, C. L.; Wood, J. A.; Malmberg, A. B.; Loas, A.; Gómez-Bombarelli, R.; Pentelute, B. L. Deep learning to design nuclear-targeting abiotic miniproteins. Nat. Chem. 2021, 13, 992–1000.

(37) Li, C.; Callahan, A. J.; Phadke, K. S.; Bellaire, B.; Farquhar, C. E.; Zhang, G.; Schissel, C. K.; Mijalis, A. J.; Hartrampf, N.; Loas, A.; Verhoeven, D. E.; Pentelute, B. L. Automated flow synthesis of peptide–PNA conjugates. ACS Cent. Sci. 2022, 8, 205–213.

(38) Dougherty, P. G.; Sahni, A.; Pei, D. Understanding cell penetration of cyclic peptides. Chem. Rev. 2019, 119, 10241–10287.

(39) Cory, A. H.; Owen, T. C.; Barlrop, J. A.; Cory, J. G. Use of an aqueous soluble tetrazolium/formazan assay for cell growth assays in culture. Cancer Commun. 1991, 3, 207–212.

(40) Vassilev, L. T.; Tovar, C.; Chen, S.; Knezevic, D.; Zhao, X.; Sun, H.; Heimbrook, D. C.; Chen, L. Selective small-molecule inhibitor reveals critical mitotic functions of human CDK1. Proc. Natl Acad. Sci. USA 2006, 103, 10660–10665.

(41) Le, J. Y.; Leonhardt, L. G.; Obeid, L. M. Cell-cycle-dependent changes in ceramide levels preceding retinoblastoma protein dephosphorylation in G2/M. Biochem. J. 1998, 334, 457–461.

(42) Ruvolo, P. P. Ceramide regulates cellular homeostasis via diverse stress signaling pathways. Leukemia 2001, 15, 1153–1160.

(43) Espaillat, M. P.; Shamseddine, A. A.; Adada, M. M.; Hannun, Y. A.; Obeid, L. M. Ceramide and sphingosine-1-phosphate in cancer, two faces of the sphinx. Transl. Cancer Res. 2015, 4, 484–499.

(44) Duchardt, F.; Fotin-Mleczek, M.; Schwarz, H.; Fischer, R.; Brock, R. A comprehensive model for the cellular uptake of cationic cell-penetrating peptides. Traffic 2007, 8, 848–866.

(45) Verdurmen, W. P. R.; Thanos, M.; Ruttekolk, I. R.; Gulbins, E.; Brock, R. Cationic cell-penetrating peptides induce ceramide formation via acid sphingomyelinase: Implications for uptake. J. Control. Release 2010, 147, 171–179.

(46) Qian, Z.; Larochelle, J. R.; Jiang, B.; Lian, W.; Hard, R. L.; Selner, N. G.; Luechapanichkul, R.; Barrios, A. M.; Pei, D. Early endosomal escape of a cyclic cell-penetrating peptide allows effective cytosolic cargo delivery. Biochemistry 2014, 53, 4034–4046.

(47) Sahni, A.; Qian, Z.; Pei, D. Cell-penetrating peptides escape the endosome by inducing vesicle budding and collapse. ACS Chem. Biol. 2020, 15, 2485–2492.

(48) Schneider, A. F. L.; Kithil, M.; Cardoso, M. C.; Lehmann, M.; Hackenberger, C. P. R. Cellular uptake of large biomolecules enabled by cell-surface-reactive cell-penetrating peptide additives. Nat. Chem. 2021, 13, 530–539.

(49) Horbay, R.; Hamraghani, A.; Ermini, L.; Holcik, S.; Beug, S. T.; Yeganeh, B. Role of ceramides and lysosomes in extracellular vesicle biogenesis, cargo sorting and release. Int. J. Mol. Sci. 2022, 23.

(50) Saha, A.; Mandal, S.; Arafiles, J. V. V.; Gómez-González, J.; Hackenberger, C. P. R.; Brik, A. Structure–uptake relationship study of DABCYL derivatives linked to cyclic cell-penetrating peptides for live-cell delivery of synthetic proteins. Angew. Chem., Int. Ed. 2022, 61, e202207551.

(51) Kim, J. A.; Åberg, C.; Salvati, A.; Dawson, K. A. Role of cell cycle on the cellular uptake and dilution of nanoparticles in a cell population. Nat. Nanotechnol. 2012, 7, 62–68.

