## Supporting Information for "Endosomolytic Peptides Enable the Cellular Delivery of Peptide Nucleic Acids"

JoLynn B. Giancola and Ronald T. Raines*
Department of Chemistry and Koch Institute for Integrative Cancer Research at MIT, Massachusetts Institute of Technology, Cambridge, Massachusetts 02139, United States

*

### I. Abbreviations Used

| | |
|---|---|
| ACN | acetonitrile |
| CHCA | α-cyano-4-hydroxycinnamic acid |
| DIC | *N*,*N*′-diisopropylcarbodiimide |
| DIPEA | *N*,*N*′-diisopropylethylamine |
| DCM | dichloromethane |
| DMEM | Dulbecco's modified Eagle's medium |
| DMF | dimethylformamide |
| DMSO | dimethyl sulfoxide |
| DTT | dithiothreitol |
| DPBS | Dulbecco's phosphate-buffered saline |
| EDTA | ethylenediaminetetraacetic acid |
| FBS | fetal bovine serum |
| Fmoc | fluorenylmethyloxycarbonyl |
| HATU | (1-[bis(dimethylamino)methylene]-1*H*-1,2,3-triazolo[4,5-b]pyridinium 3-oxid hexafluorophosphate) |
| MALDI–TOF | matrix-assisted laser desorption/ionization time-of-flight |
| MFI | median fluorescence intensity |
| MS | mass spectrometry |
| PEG | polyethylene glycol |
| Q–TOF | quadrupole time-of-flight |
| RP-HPLC | reversed-phase high-performance liquid chromatography |
| RPM | revolutions per minute |
| SPPS | solid-phase peptide synthesis |
| TFA | trifluoroacetic acid |
| TIS | triisopropylsilane |

### II. General Methods

*Reagents and Solvents*. Commercially available reagents and solvents were reagent grade or better, and were used directly without further purification. Reagents and solvents were from Sigma–Aldrich (St. Louis, MO) unless otherwise specified. Amino acids were from Chem-Impex International (Wood Dale, IL). Rink Amide ProTide resin (LL) and Oxyma Pure were from CEM (Matthews, NC). DIC and 4-methylpiperidine were from Oakwood Chemical (Tampa, FL). Anhydrous DMSO, TFA, and TIS were from Sigma–Aldrich (St. Louis, MO). L17E was from BioSynth (Staad, Switzerland). PNA654 ((O-GCTATTACCTTAACCCAG-Lys(C6SH) and (O-GCTATTACCTTAACCCAG-Lys(DBCO)) were from PNA Bio (Thousand Oaks, CA). Water was obtained from a Milli-Q IQ 7000 purification system from MilliporeSigma (Burlington, MA) and had a resistivity of $18.2 \times 10^6$ Ω cm.

*Peptide Synthesis*. All amino acids used in SPPS were of L stereochemistry and were protected at their N terminus with Fmoc from Chem-Impex. Peptides were synthesized on Rink Amide ProTide Resin (LL) (0.05 or 0.1 mmol, 0.59 mmol/g, 1.0 equiv) with a Liberty Blue Automated Microwave Peptide Synthesizer from CEM following the manufacturer's standard procedures. Standard solutions of Oxyma Pure (1.0 M in DMF), DIC (0.5 M in DMF), 4-methylpiperidine

(20% v/v in DMF), and Fmoc-protected amino acids (0.2 M in DMF) were used in coupling and deprotection steps. Peptides were purified with a 1260 Infinity II Preparative LC System from Agilent Technologies equipped with a XSelect Peptide CSH C18 OBD prep column (130 Å pore size, 5 µm particle size, 19 mm × 250 mm of width × length) from Waters (Milford, MA).

*Mass Spectrometry.* Mass spectra of small molecules were acquired on an LCT electrospray ionization (ESI) 1260 Infinity II instrument from Agilent Technologies (Santa Clara, CA) and an LC–MS column (Agilent Technologies, Poroshell 120, SB C18-reversed-phase, length 50 mm, internal diameter: 2.1 mm, particle size: 2.7 micron) with a gradient of 10−95% v/v MeCN (0.1% v/v formic acid) in water (0.1% v/v formic acid) over 10 min. To minimize the fragmentation of diazo moieties, the MSD parameters were set as follows: capillary voltage, 3000 V; drying gas temperature, 350 °C; gas flow, 13/min; fragmentor voltage, 30 V; nebulizer pressure, 35 psig; and cycle time, 0.83 s/cycle. HRMS of peptides and small molecules was performed with Agilent 6545 Q–TOF mass spectrometer coupled to an Agilent Infinity 1260 LC system (Q–TOF). The crude molecular mass of peptides was determined on α-cyano-4-hydroxycinnamic acid matrix by matrix-assisted laser desorption/ionization time-of-flight (MALDI–TOF) mass spectrometry with a microflex LRF instrument from Bruker (Billerica, MA). MALDI samples were all desalted using DOWEX 50WX4-400 strong cation-exchange resin (CAS 11113-61-4) before spotting 1:1 v/v with the appropriate matrix. A more accurate assessment of the molecular mass of peptides and conjugates was carried out using ESI mass spectrometry on a 6530C Accurate-Mass Q–TOF MS equipped with a Poroshell 120 column (EC-C18 2.7 micron, 3.0 × 150 mm) (product #693975-302T) from Agilent Technologies. A gradient of 5–40% v/v MeCN (0.1% v/v formic acid) in water (0.1% v/v formic acid) over 16 min was used unless otherwise indicated. Before Q–TOF LC–MS analysis, all samples were passed through a Spin-X Centrifugal Tube Filter (0.22-µm, cellulose acetate membrane) from R&D Systems (Minneapolis, MN).

*Compound Purity.* The purity of small molecules was judged to be ≥95%, as assessed by mass spectrometry and RP-HPLC using an LCsdMS column and gradient of 10−95% v/v MeCN (0.1% v/v formic acid) in water (0.1% v/v formic acid) over 10 min unless indicated otherwise. The purity of peptides was accessed with a 1260 Infinity II Preparative LC System from Agilent Technologies equipped with an EC NUCLEOSIL 100-5 C18 analytical column (100 Å, 5 µm, 4.6 mm × 250 mm) from Macherey-Nagel (Düren, Germany) or 1200 Infinity System from Agilent Technologies equipped with a Microsorb-MV 100-5 C18 column (100 Å, 5 µm, 4.6 mm × 250 mm) from Varian (Palo Alto, CA).

*Biological Reagents, Supplies, and Instrumentation.* Titer Plate Shaker was from Labline Instruments (Melrose Park, IL). TAMRA-labelled peptide concentrations were determined with a DS-11 UV–vis spectrophotometer from DeNovix (Wilmington, DE). Pierce dye-removal columns were from ThermoFisher Scientific (product #22858). Amicon Ultra 0.5-mL 10K MWCO centrifugal filter units were from MilliporeSigma (product #UFC501024). MTS readings were collected with a Spark plate reader from Tecan (Männedorf, Switzerland). DMEM (product #11995065) was from ThermoFisher Scientific. FBS (product #45001-108) was from Corning (Corning, NY). Penicillin-streptomycin (10,000 U/mL) (product #15140122) was from ThermoFisher Scientific. Trypsin–EDTA (0.25% w/v) with phenol red was from ThermoFisher Scientific (product #25200056). Cells were counted using a Countess II FL automated cell counter (product #AMQAF1000) with Countess Cell Counting Chamber Slides (product #C10283) from ThermoFisher Scientific. Differential interference contrast and epifluorescent live cells images were acquired using an epifluorescent EVOS M7000 Imaging System (product #AMF7000) from ThermoFisher Scientific. IbiTreat 18-well plates (product #81816) were from Ibidi (Fitchburg,

WI). DPBS with $Ca^{2+}/Mg^{2+}$ (product #14040141) and DPBS without $Ca^{2+}/Mg^{2+}$ (product #14190144) were from Gibco (Waltham, MA). FluoroBrite DMEM (product #A1896701) was from ThermoFisher Scientific. Life Technologies Attune NxT flow cytometer, SYTOX Red Dead-Cell Indicator (product #S34859) and SYTOX Blue Dead-Cell Indicator (product #S34857) from ThermoFisher Scientific were used for flow cytometry.

*Conditions*. All procedures were performed in air at ambient temperature (~22 °C) and pressure (1.0 atm) unless indicated otherwise.

#### III. Peptide Synthesis and Characterization

L17E- and L17ER$_4$-containing peptides were prepared by fusing the same general procedure. R10-containing peptides were prepared under different conditions depending on the functional groups present in the peptide. Synthesis methods for R10-containing peptides are provided before the analytical characterization is given, and sequences are provided in Table S1.

##### General Synthesis Method for L17E and L17ER$_4$ Peptides

*Synthesis*. Linear peptides with sequences defined in Table S1 were synthesized with single-couplings of amino acid monomers. When applicable, Fmoc-Cys(Trt)-OH, Fmoc-Lys(N$_3$)-OH, and Fmoc-His(Boc)-OH were coupled for 10 min at 50 °C under standard microwave conditions. After the synthesis, the resin was transferred to a 24-mL polypropylene luer-lock syringe equipped with a filter frit.

*Cleavage and Precipitation*. Peptides were first washed with DMF (5×) followed by DCM (15×) and were cleaved from the resin over 3 h using cleavage cocktail (82.5% v/v TFA, 5% w/v phenol, 5% v/v $H_2O$, 5% v/v thioanisole, 2.5% v/v 3,6-dioxa-1,8-octanedithiol) at 2× the volume of DCM-swelled resin. The resin was washed with an additional 4 mL of cleavage cocktail, and the pooled cleavage eluate was blown under a stream of $N_2$(g) or Ar(g) to evaporate the cleavage cocktail. When the peptide had concentrated to a thick orange oil, the peptide was precipitated in 50 mL of ice-cold diethyl ether. The peptide was pelleted by centrifugation for 10 min at 1500 RPM at 4 °C. The ether supernatant was decanted, and the crude peptide was stored at −70 °C until purified by RP-HPLC.

*Purification*. RP-HPLC was performed by using a VP 250/21 Nucleosil 100-5 C18 column from Macherey–Nagel (Bethlehem, PA) and a 1260 Infinity II instrument from Agilent Technologies (Santa Clara, CA). The crude peptide was reconstituted in a minimal amount of ACN, passed through a 0.22-μm PFTE filter, and separated using a gradient of 15–40% v/v ACN in $H_2O$ containing TFA (0.1% v/v) over 40 min. Fraction purity was assessed by MALDI–TOF MS in positive mode using a microflex LRF instrument from Bruker (Billerica, MA) and a CHCA matrix or by LC–MS. Pure fractions were pooled and lyophilized using a FreeZone benchtop instrument from Labconco (Kansas City, MO) to yield the peptides as fluffy white TFA salts. Final purity was assessed by RP-HPLC using an EC 250/4.6 Nucleosil 100-5 C18 column from Macherey–Nagel and a 1260 Infinity II instrument from Agilent Technologies. Synthesis scale, yields, and analytical characterization are described below.

**Table S1. Peptide Sequences**

| Peptide | Sequence |
|---|---|
| L17E | H-IWLTALKFLGKHAAKHEAKQQLSKL-amide |
| L17E-GGK($N_3$) | H-IWLTALKFLGKHAAKHEAKQQLSKLGGK($N_3$)-amide |
| K($N_3$)GG-L17E | H-K($N_3$)GGIWLTALKFLGKHAAKHEAKQQLSKL-amide |
| $L17ER_4$ | H-IWLTALKFLGKHAAKHEAKQQLSKLGRRRR-amide |
| $L17ER_4$-GGK($N_3$) | H-IWLTALKFLGKHAAKHEAKQQLSKLGRRRRGGK($N_3$)-amide |
| K($N_3$)GG-$L17ER_4$ | H-K($N_3$)GGIWLTALKFLGKHAAKHEAKQQLSKLGRRRR-amide |
| R10 | H-RRRRRRRRRR-amide |
| Azide-R10 | $N_3$-$PEG_2$-RRRRRRRRRR-amide |

#### Analytical Characterization of L17E- and $L17ER_4$-containing Peptides

### L17E

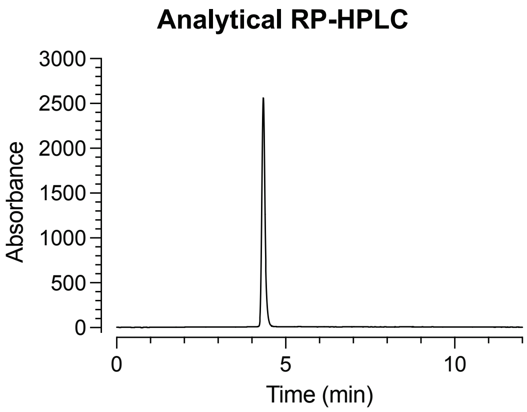

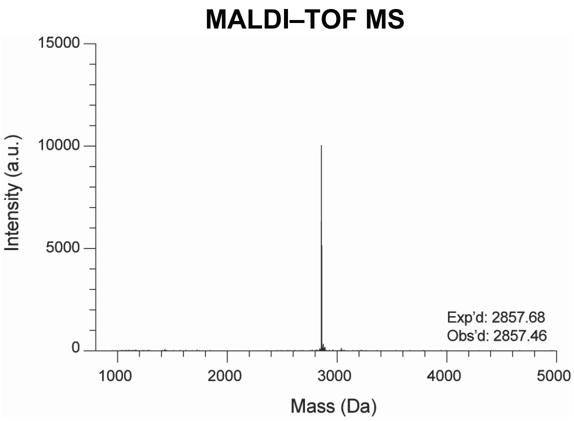

##### Yield of L17E
L17E was synthesized on a 0.1 mmol scale, and the product was isolated via RP-HPLC to yield a trifluoroacetate salt of L17E (94.9 mg, 26.8% yield; expected molecular mass of the L17E·$TFA_6$ salt, 3543.59).

##### Analytical RP-HPLC of L17E
Purity was assessed with analytical RP-HPLC using a linear gradient of 10–70% v/v acetonitrile in water containing TFA (0.1% v/v) at 2 mL/min over 10 min ($\lambda$ = 218 nm). Product was assessed as being >99% pure by analytical RP-HPLC.

##### LC/MS Analysis of L17E
Exp'd $m/z$ $[M + 5H]^{5+}$, 572.5; $[M + 6H]^{6+}$, 477.3; $[M + 7H]^{7+}$, 409.2
Obs'd $m/z$ $[M + 5H]^{5+}$, 572.8; $[M + 6H]^{6+}$, 477.5; $[M + 7H]^{7+}$, 409.5

#### L17E-GGK($N_3$)

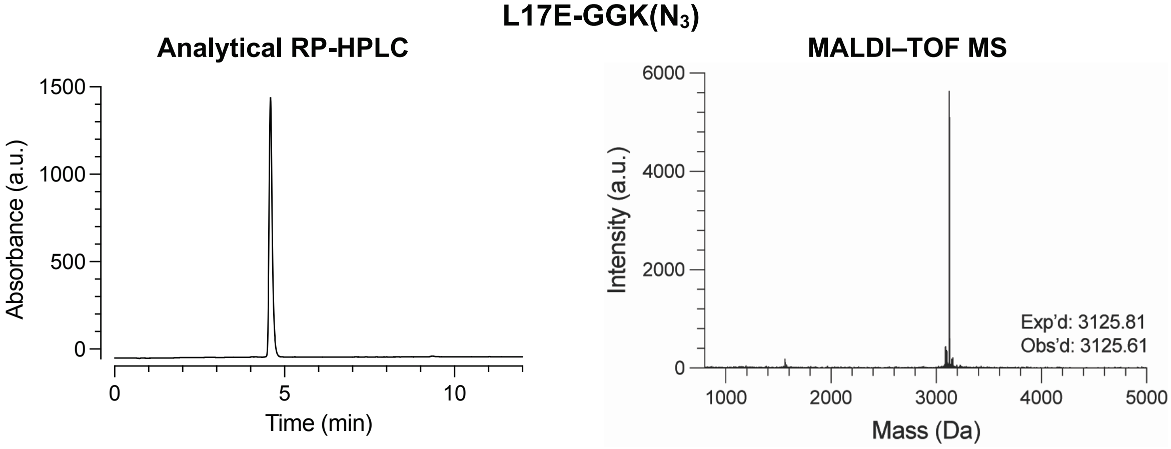

##### Yield of L17E-GGK($N_3$)

L17E-GGK($N_3$) was synthesized on a 0.05 mmol scale, and the product was isolated via RP-HPLC to yield a trifluoroacetate salt of L17E-GGK($N_3$) (39.8 mg, 20.9% yield, expected molecular mass of the L17E-GGK($N_3$)·$TFA_6$ salt, 3811.87).

##### Analytical RP-HPLC of L17E-GGK($N_3$)

Purity was assessed with analytical RP-HPLC using a linear gradient of 10–70% v/v acetonitrile in water containing TFA (0.1% v/v) at 2 mL/min over 12 min ($\lambda$ = 218 nm). Product was assessed as being >99% pure by analytical RP-HPLC.

##### LC/MS Analysis of L17E-GGK($N_3$)

Exp'd *m/z* $[M + 3H]^{3+}$, 1042.9; $[M + 4H]^{4+}$, 782.5; $[M + 5H]^{5+}$, 626.2; $[M + 6H]^{6+}$, 522.0; $[M + 7H]^{7+}$, 447.5
Obs'd *m/z* $[M + 3H]^{3+}$, 1043.3; $[M + 4H]^{4+}$, 782.8; $[M + 5H]^{5+}$, 626.5; $[M + 6H]^{6+}$, 522.2; $[M + 7H]^{7+}$, 447.7

## K($N_3$)GG-L17E

K($N_3$)GG-L17E

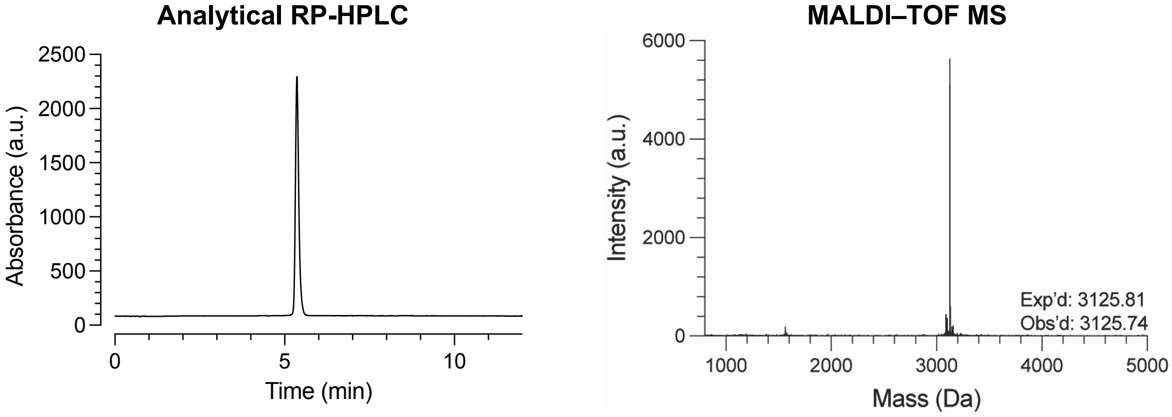

##### Yield of K($N_3$)GG-L17E

K($N_3$)GG-L17E was synthesized on a 0.05 mmol scale, and the product was isolated via RP-HPLC to yield a trifluoroacetate salt of K($N_3$)GG-L17E (39.2 mg, 20.6% yield; expected molecular mass of the K($N_3$)GG-L17E·$TFA_6$ salt, 3811.87).

##### Analytical RP-HPLC of K($N_3$)GG-L17E

Purity was assessed with analytical RP-HPLC using a linear gradient of 10–70% v/v acetonitrile in water containing TFA (0.1% v/v) at 2 mL/min over 12 min ($\lambda$ = 218 nm). Product was assessed as being >99% pure by analytical RP-HPLC.

##### LC/MS Analysis of K($N_3$)GG-L17E

Exp'd *m*/*z* $[M + 5H]^{5+}$, 626.2; $[M + 6H]^{6+}$, 522.0; $[M + 7H]^{7+}$, 447.5
Obs'd *m*/*z* $[M + 5H]^{5+}$, 626.5; $[M + 6H]^{6+}$, 522.2; $[M + 7H]^{7+}$, 447.8

## L17ER$_4$

L17ER$_4$

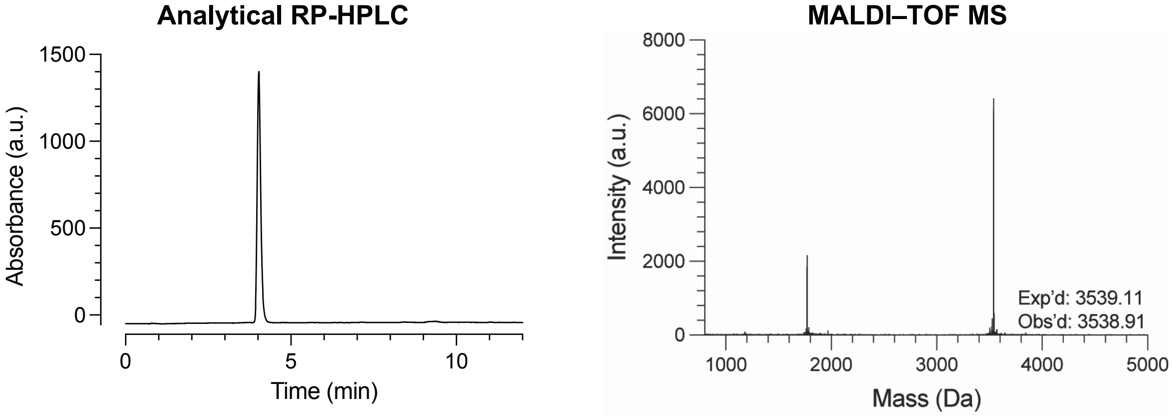

##### Yield of L17ER$_4$

L17ER$_4$ was synthesized on a 0.1 mmol scale, and the product was isolated via RP-HPLC to yield a trifluoroacetate salt of L17ER$_4$ (45.9 mg, 9.8% yield; expected molecular mass of the L17ER$_4$·TFA$_{10}$ salt, 4681.48).

##### Analytical RP-HPLC of L17ER$_4$

Purity was assessed with analytical RP-HPLC using a linear gradient of 10–70% v/v acetonitrile in water containing TFA (0.1% v/v) at 2 mL/min over 12 min ($\lambda$ = 218 nm). Product was assessed as being >98% pure by analytical RP-HPLC.

##### LC/MS Analysis of L17ER$_4$

Exp'd *m/z* $[M + 5H]^{5+}$, 708.8; $[M + 6H]^{6+}$, 590.9; $[M + 7H]^{7+}$, 506.6; $[M + 8H]^{8+}$, 443.4; $[M + 9H]^{9+}$, 394.2
Obs'd *m/z* $[M + 5H]^{5+}$, 709.2; $[M + 6H]^{6+}$, 591.1; $[M + 7H]^{7+}$, 506.8; $[M + 8H]^{8+}$, 443.5; $[M + 9H]^{9+}$, 394.4

#### L17ER$_4$-GGK(N$_3$)

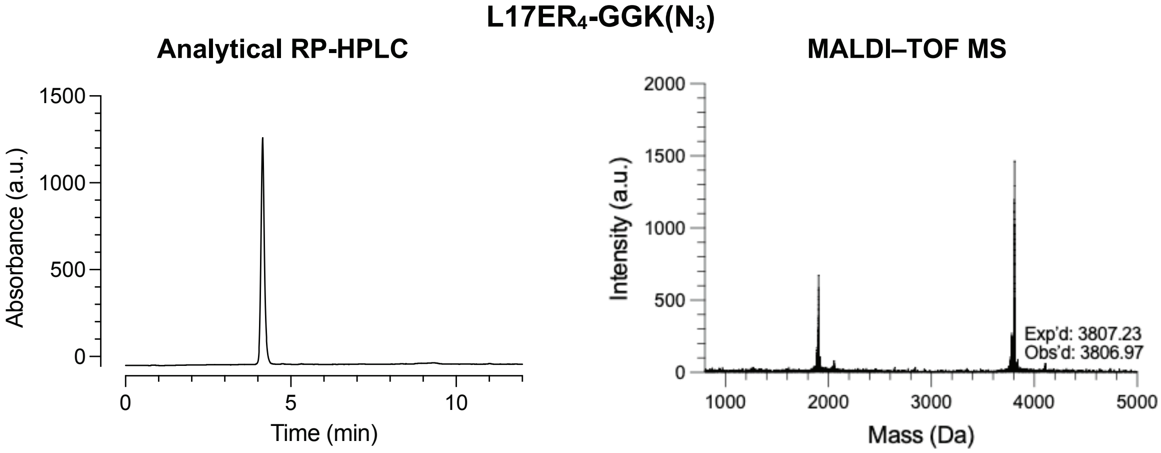

**Yield of L17ER$_4$-GGK(N$_3$)**

L17ER$_4$-GGK(N$_3$) was synthesized on a 0.05 mmol scale, and the product was isolated via RP-HPLC to yield a trifluoroacetate salt of L17ER$_4$-GGK(N$_3$) (34.4 mg, 13.9% yield, expected molecular mass of the L17ER$_4$-GGK(N$_3$)·TFA$_{10}$ salt, 4949.75).

**Analytical RP-HPLC of L17ER$_4$-GGK(N$_3$)**

Purity was assessed with analytical RP-HPLC using a linear gradient of 10–70% v/v acetonitrile in water containing TFA (0.1% v/v) at 2 mL/min over 12 min ($\lambda$ = 218 nm). Product was assessed as being >98% pure by analytical RP-HPLC.

**LC/MS Analysis of L17ER$_4$-GGK(N$_3$)**

Exp'd *m*/*z* $[M + 5H]^{5+}$, 762.4; $[M + 6H]^{6+}$, 635.5; $[M + 7H]^{7+}$, 544.9; $[M + 8H]^{8+}$, 476.9; $[M + 9H]^{9+}$, 424.0

Obs'd *m*/*z* $[M + 5H]^{5+}$, 762.8; $[M + 6H]^{6+}$, 635.8; $[M + 7H]^{7+}$, 545.2; $[M + 8H]^{8+}$, 477.1; $[M + 9H]^{9+}$, 424.2

## K($N_3$)GG-L17E$R_4$

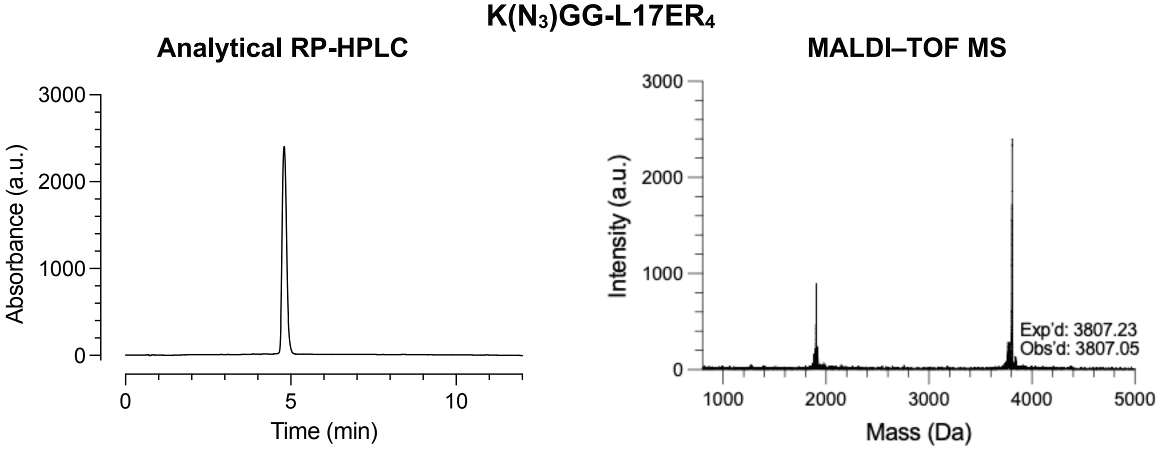

##### Yield of K($N_3$)GG-L17E$R_4$

K($N_3$)GG-L17E$R_4$ was synthesized on a 0.05 mmol scale, and the product was isolated via RP-HPLC to yield a trifluoroacetate salt of K($N_3$)GG-L17E$R_4$ (22.6 mg, 9.1% yield, expected molecular mass of the K($N_3$)GG-L17E$R_4$·TF$A_{10}$ salt, 4949.75).

##### Analytical RP-HPLC of K($N_3$)GG-L17E$R_4$

Purity was assessed with analytical RP-HPLC using a linear gradient of 10–70% v/v acetonitrile in water containing TFA (0.1% v/v) at 2 mL/min over 12 min ($\lambda$ = 218 nm). Product was assessed as being >99% pure by analytical RP-HPLC.

##### LC/MS Analysis of K($N_3$)GG-L17E$R_4$

Exp'd *m*/*z* $[M + 5H]^{5+}$, 762.4; $[M + 6H]^{6+}$, 635.5; $[M + 7H]^{7+}$, 544.9; $[M + 8H]^{8+}$, 476.9; $[M + 9H]^{9+}$, 424.0; $[M + 10H]^{10+}$, 381.7
Obs'd *m*/*z* $[M + 5H]^{5+}$, 762.8; $[M + 6H]^{6+}$, 635.8; $[M + 7H]^{7+}$, 545.1; $[M + 8H]^{8+}$, 477.1; $[M + 9H]^{9+}$, 424.2; $[M + 10H]^{10+}$, 381.9

**R10**

*Synthesis*. A linear R10 peptide with the sequence Fmoc-(Arg(Pbf))$_{10}$–$NH_2$ was synthesized with double-couplings of amino acid monomers. After the synthesis, the resin was transferred to a 24-mL polypropylene luer-lock syringe equipped with a filter frit.

*Cleavage and Precipitation*. Peptides were first washed with DMF (5×) followed by DCM (15×) and were cleaved from the resin over 5 h (7 mL mixture of 95% v/v TFA, 2.5% v/v , 2.5% v/v DTT). The resin was washed with an additional 4 mL of cleavage cocktail, and the pooled cleavage eluate was blown under a stream of $N_2$(g) to evaporate the cleavage cocktail. When the peptide had concentrated to a thick red oil, the peptide was precipitated in 50 mL of ice-cold diethyl ether. The peptide was pelleted by centrifugation for 10 min at 1500 RPM at 4 °C. The ether supernatant was decanted, and the crude peptide was stored at −70 °C until purified by RP-HPLC.

*Purification*. RP-HPLC was performed by using a VP 250/21 Nucleosil 100-5 C18 column from Macherey–Nagel (Bethlehem, PA) and a 1260 Infinity II instrument from Agilent Technologies (Santa Clara, CA). The crude peptide was reconstituted in a minimal amount of ACN, passed through a 0.22-μm PFTE filter, and separated using a gradient of 5–35% v/v ACN in $H_2O$ containing TFA (0.1% v/v) over 35 min. Fraction purity was assessed by MALDI–TOF MS in positive mode using a microflex LRF instrument from Bruker (Billerica, MA) and a CHCA matrix. Pure fractions were pooled and lyophilized using a FreeZone benchtop instrument from Labconco (Kansas City, MO) to yield the peptide as a fluffy white TFA salt (25.2 mg, 8.55 μmol, 17.1% yield). The molecular mass of the purified material was confirmed by MALDI–TOF MS (molecular mass, 1578.92 g/mol; molecular mass TFA$_{12}$-salt, 2947.16 g/mol). Final purity was assessed by RP-HPLC using an EC 250/4.6 Nucleosil 100-5 C18 column from Macherey–Nagel and a 1260 Infinity II instrument from Agilent Technologies.

**R10**

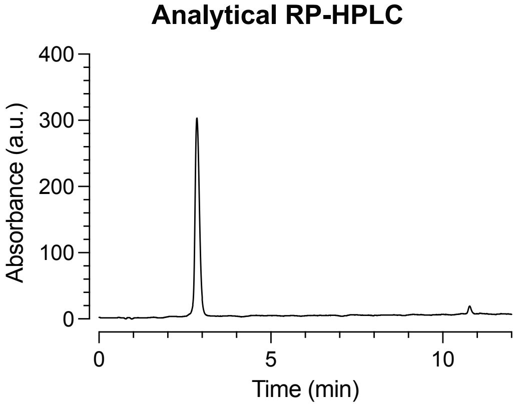

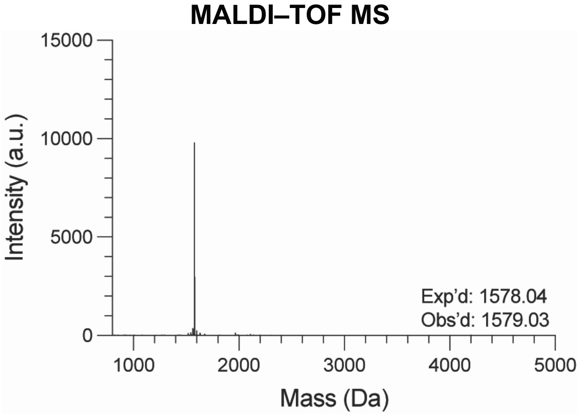

##### Yield of R10

R10 was synthesized on a 0.05 mmol scale, and the product was isolated via RP-HPLC to yield a trifluoroacetate salt of R10 (25.2 mg, 17.1% yield, expected molecular mass of the R10-SH·$TFA_{12}$ salt = 2947.16).

##### Analytical RP-HPLC of R10

Purity was assessed with analytical RP-HPLC using a linear gradient of 5–25% v/v B over 12 min at 2 mL/min ($\lambda$ = 218 nm). Product was assessed as being >96% pure by analytical RP-HPLC.

##### LC/MS Analysis of R10

Exp'd *m*/*z*: $[M + 3H]^{3+}$, 527.0; $[M + 4H]^{4+}$, 395.5; $[M + 5H]^{5+}$, 316.6; $[M + 6H]^{6+}$, 264.0
Obs'd *m*/*z*: $[M + 3H]^{3+}$: 527.1, $[M + 4H]^{4+}$: 395.6, $[M + 5H]^{5+}$: 316.7, $[M + 6H]^{6+}$: 264.1

#### Azide-R10

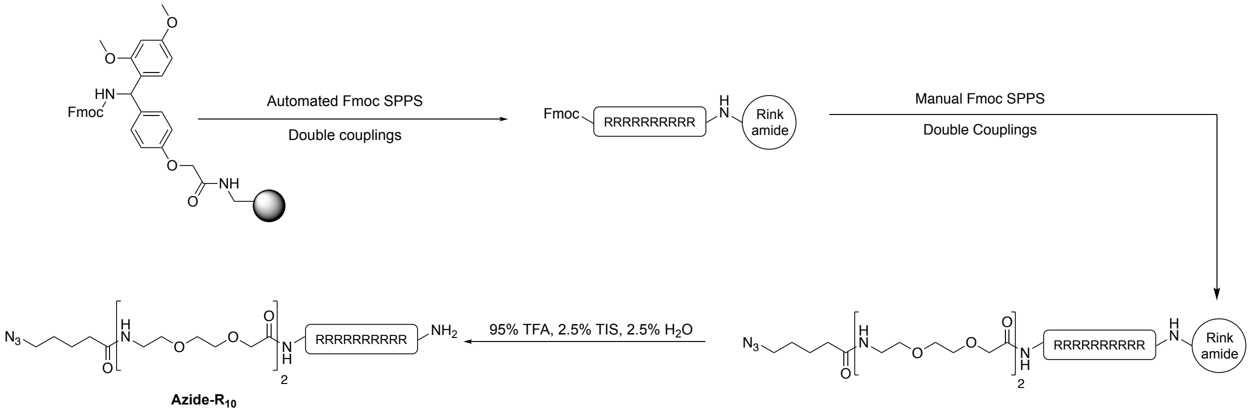

*Synthesis*. After the synthesis of the Fmoc-(Arg(Pbf))$_{10}$-$NH_2$ peptide as described above, the resin was transferred to a 24-mL polypropylene luer-lock syringe equipped with a filter frit, where further elaboration of the peptide was performed by hand according to traditional Fmoc-based methods. Fmoc-8-amino-3,6-dioxaoctanoic acid (4 equiv) was double-coupled using HATU (4 equiv) and DIPEA (8 equiv) in 3 mL of DMF. A final Fmoc deprotection was performed, and subsequent double-coupling using 5-azidopentanoic acid (5 equiv) was performed using HATU (5 equiv) and DIPEA (10 equiv) in 2 mL of DMF for 1 h. Fmoc protecting groups were cleaved by treating the resin with a solution of 20% v/v *N*-methylpiperidine in DMF (2 × 5 min each time). The resin was washed after deprotection steps and in between amino acid couplings using DMF (5×), DCM (5×), and DMF (5×).

*Cleavage and Precipitation*. Peptides were first washed with DMF (5×) followed by DCM (15×) and cleaved from the resin for 3 h (6 mL mixture of 95% v/v TFA, 2.5% v/v TIS, 2.5% v/v water). The resin was washed with an additional 4 mL of cleavage cocktail, and the pooled cleavage eluate was blown under a stream of $N_2$(g) to evaporate the cleavage cocktail. When the peptide had concentrated to a thick brown oil, the peptide was precipitated in 50 mL of ice-cold diethyl ether. The peptide was pelleted by centrifugation for 10 min at 1500 RPM at 4 °C. The ether supernatant was decanted, and the crude peptide was stored at −70 °C until purified by RP-HPLC.

*Purification*. RP-HPLC was performed using a VP 250/21 Nucleosil 100-5 C18 column from Macherey–Nagel (Bethlehem, PA) and a 1260 Infinity II instrument from Agilent Technologies (Santa Clara, CA). The crude peptide was reconstituted in a minimal amount of ACN, passed through a 0.22 µm PFTE filter, and separated using a gradient of 5–35% ACN in $H_2O$ containing TFA (0.1% v/v) over 35 min. Fraction purity was assessed by MALDI–TOF MS in positive mode using a microflex LRF instrument from Bruker (Billerica, MA) and a CHCA matrix. Pure fractions were pooled and lyophilized using a FreeZone benchtop instrument from Labconco (Kansas City, MO) to yield the peptide as a fluffy white TFA salt (17.8 mg, 5.48 µmol, 5.5%). The molecular mass of the purified material was confirmed by MALDI–TOF MS (molecular mass, 1994.37 g/mol; molecular mass TFA$_{11}$-salt, 3248.63 g/mol). Final purity was assessed by RP-HPLC using an EC 250/4.6 Nucleosil 100-5 C18 column from Macherey–Nagel and a 1260 Infinity II instrument from Agilent Technologies.

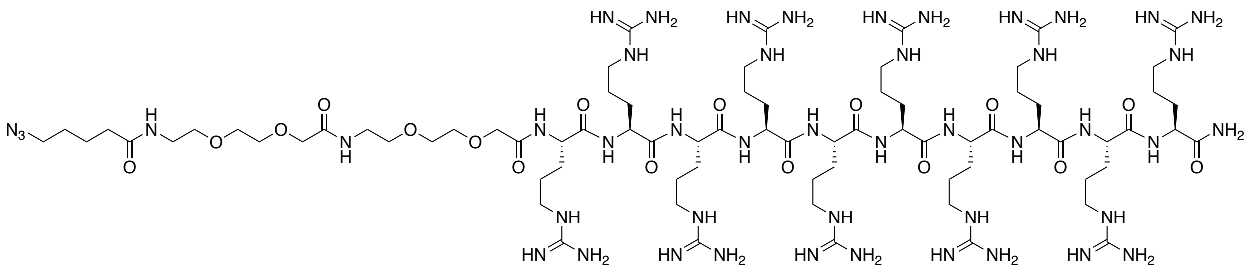

Exact Mass: 1993.24
Molecular Weight: 1994.37

**Analytical RP-HPLC**

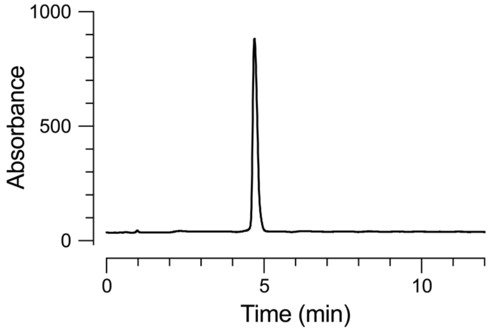

**MALDI–TOF MS**

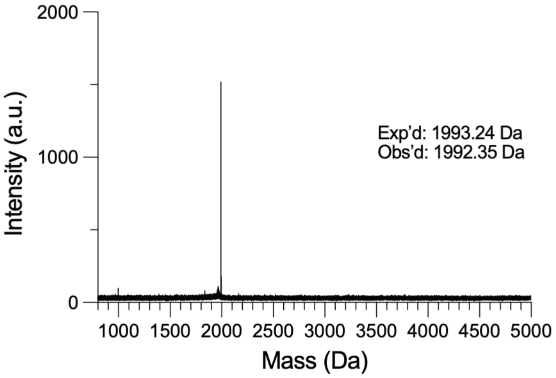

##### Yield of Azide-R10

Azide-R10 was synthesized on a 0.05 mmol scale, and the product was isolated via RP-HPLC to yield a trifluoroacetate salt of Azide-R10 (17.8 mg, 5.5% yield, expected molecular mass of the R10-SH·$TFA_{11}$ salt, 3248.63).

##### Analytical RP-HPLC of Azide-R10

Purity was assessed with analytical RP-HPLC using a linear gradient of 5–25% v/v B at 2 mL/min over 12 min ($\lambda$ = 210 nm). Product was assessed as being >98% pure by analytical HPLC.

##### LC/MS Analysis of Azide-R10

Exp'd *m/z*: $[M + 3H]^{3+}$, 665.4; $[M + 4H]^{4+}$, 499.3; $[M + 5H]^{5+}$, 399.6; $[M + 6H]^{6+}$, 333.2
Obs'd *m/z* $[M + 3H]^{3+}$, 655.5; $[M + 4H]^{4+}$, 499.5; $[M + 5H]^{5+}$, 399.8; $[M + 6H]^{6+}$, 333.1

### IV. Peptide–Fluorophore Conjugate Synthesis

#### General Procedure

To conjugate the azide-containing peptides to TAMRA-DBCO (product #40694) from BroadPharm (San Diego, CA), a 1 mM stock solution of the azide-containing peptide and a 5 mM stock soluv/v tion of the TAMRA-DBCO were prepared in water. The azide-containing peptide (1 equiv) and TAMRA-DBCO (1.1 equiv) were combined water. The reaction mixture was incubated for 48 h under agitation, and reaction progress was monitored by Q–TOF MS. When the unconjugated azide-containing peptide was consumed, excess dye was removed using peptide desalting columns (product #89852) from ThermoFisher Scientific according to the manufacturer's recommendations, with a 40% v/v ACN/$H_2O$ with 0.1% v/v TFA eluant. Peptide-fluorophore conjugates were pooled into a microcentrifuge tube, lyophilized, and reconstituted in water. Peptide-fluorophore conjugate concentrations were assessed by absorbance values of the TAMRA dye with a DeNovix DS-11 spectrophotometer.

#### L17E–TAMRA

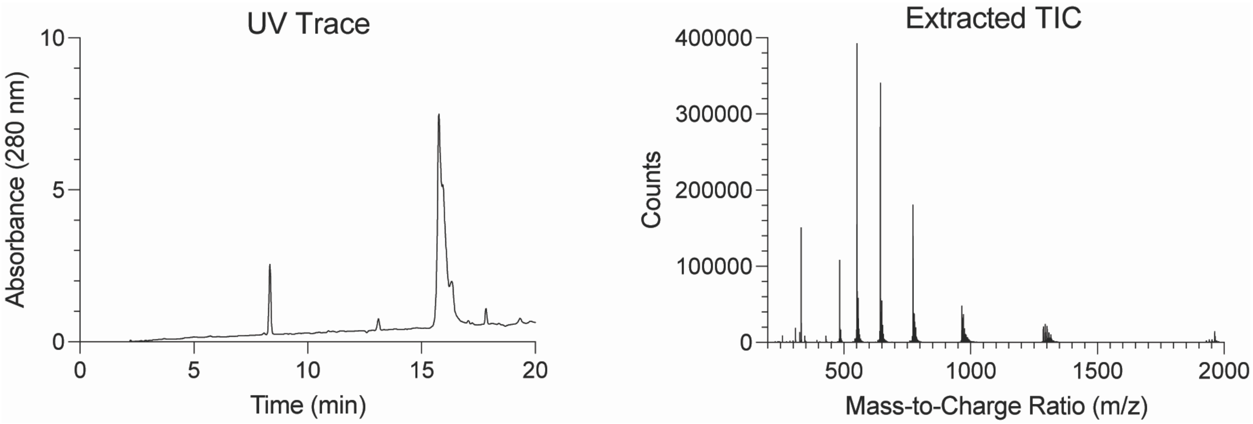

| | | |
|---|---|---|
| L17E–TAMRA | Exp'd mass | 3858.61 |
| | Obs'd mass | 3858.19 |

**Figure S1.** Representative deconvoluted Q–TOF mass spectra of L17E–TAMRA.

#### L17ER$_4$–TAMRA

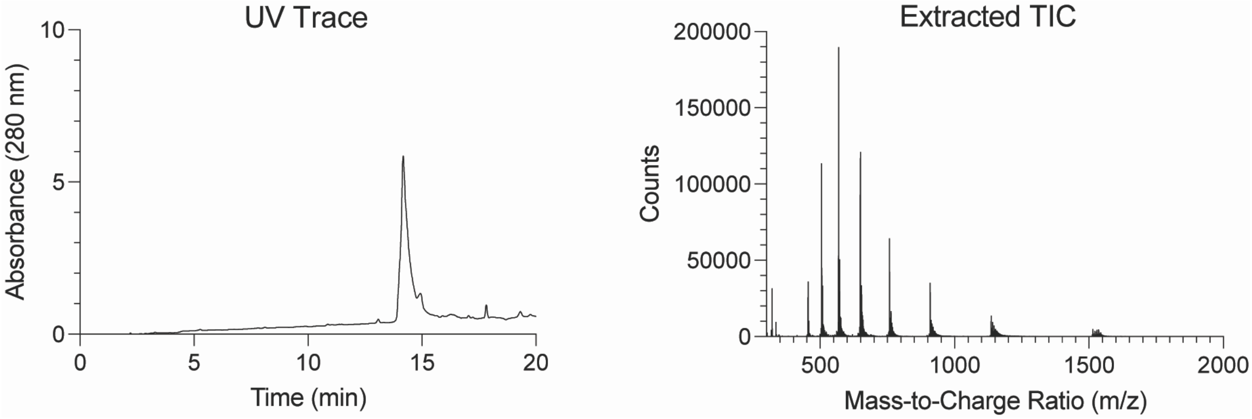

| | | |
|---|---|---|
| L17ER$_4$–TAMRA | Exp'd mass | 4540.42 |
| | Obs'd mass | 4540.62 |

**Figure S2.** Representative deconvoluted Q–TOF mass spectra of L17ER$_4$–TAMRA.

**TAMRA–L17E**

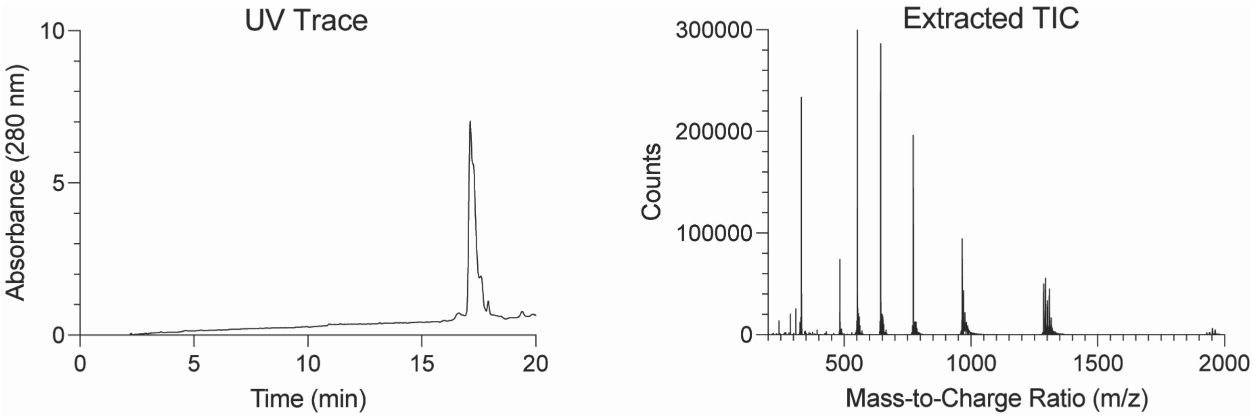

| | | |
|---|---|---|
| TAMRA–L17E | Exp'd mass | 3858.61 |
| | Obs'd mass | 3858.19 |

**Figure S3.** Representative deconvoluted Q–TOF mass spectra of TAMRA–L17E.

**TAMRA–L17ER$_4$**

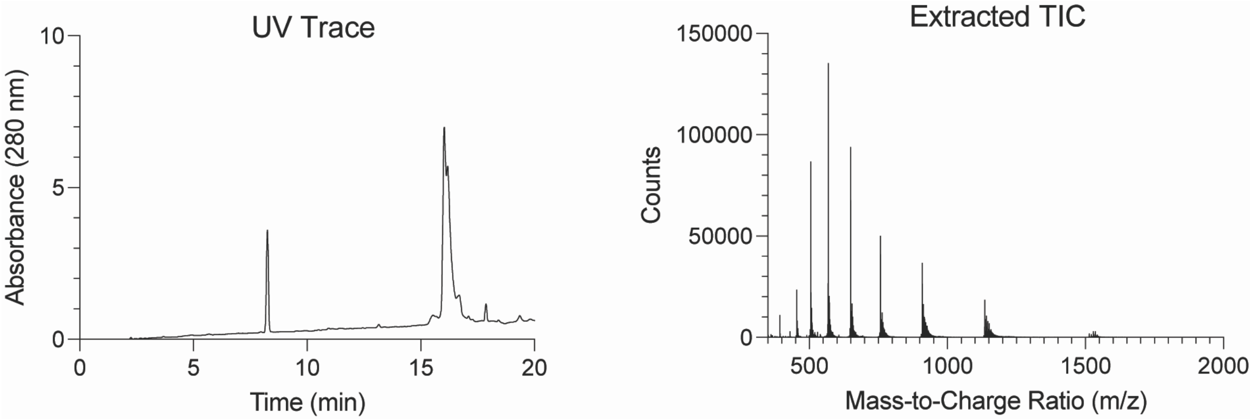

| | | |
|---|---|---|
| TAMRA–L17ER$_4$ | Exp'd mass | 4540.42 |
| | Obs'd mass | 4540.62 |

**Figure S4.** Representative deconvoluted Q–TOF mass spectra of TAMRA–L17ER$_4$.

#### V. Peptide–PNA Conjugate Synthesis

##### L17E–PNA

To conjugate the azide-containing L17E-GGK($N_3$) peptide to PNA-DBCO, a 1 mM stock solution of the L17E–azide peptide and a 1 mM stock solution of the PNA-DBCO were prepared in water. The PNA and peptide were combined stoichiometrically in water. The reaction mixture was incubated for 72 h at 4 °C under agitation, and reaction progress was monitored by Q–TOF MS. The product was used directly without further purification. The peak at RT = 9 min corresponds to hydrolyzed PNA-DBCO.[1, 2]

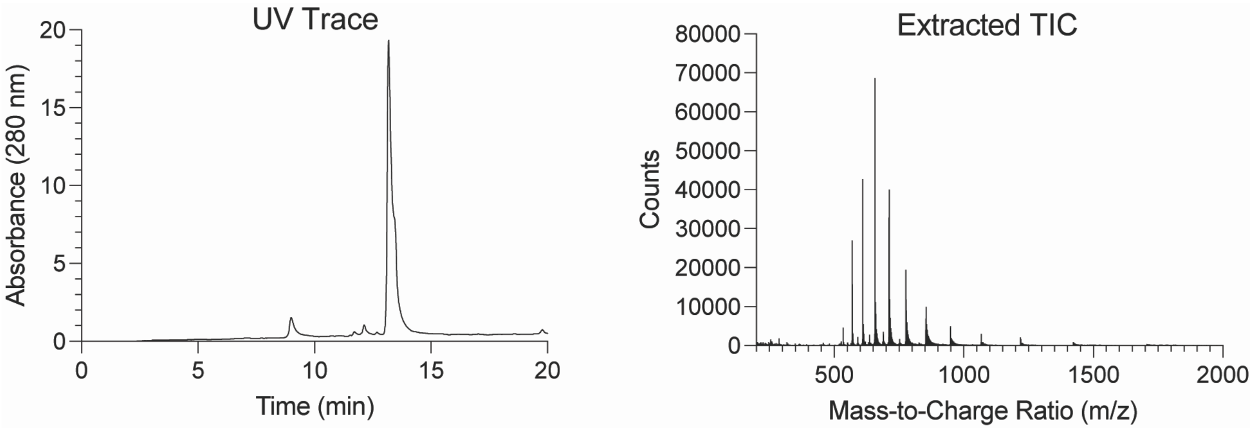

| | | |
|---|---|---|
| L17E–PNA | Exp'd mass | 8532.05 |
| | Obs'd mass | 8531.28 |

**Figure S5.** Representative deconvoluted Q–TOF mass spectra of L17E–PNA.

#### $L17ER_4$–PNA

To conjugate the azide-containing $L17ER_4$-GGK($N_3$) peptide to PNA-DBCO, a 1 mM stock solution of the $L17ER_4$–azide peptide and a 1 mM stock solution of the PNA-DBCO were prepared in water. The PNA and peptide were combined stoichiometrically in water. The reaction mixture was incubated for 72 h at 4 °C under agitation, and reaction progress was monitored by Q–TOF MS. The product was used directly without further purification. The peak at RT = 9 min corresponds to hydrolyzed PNA-DBCO.[1, 2]

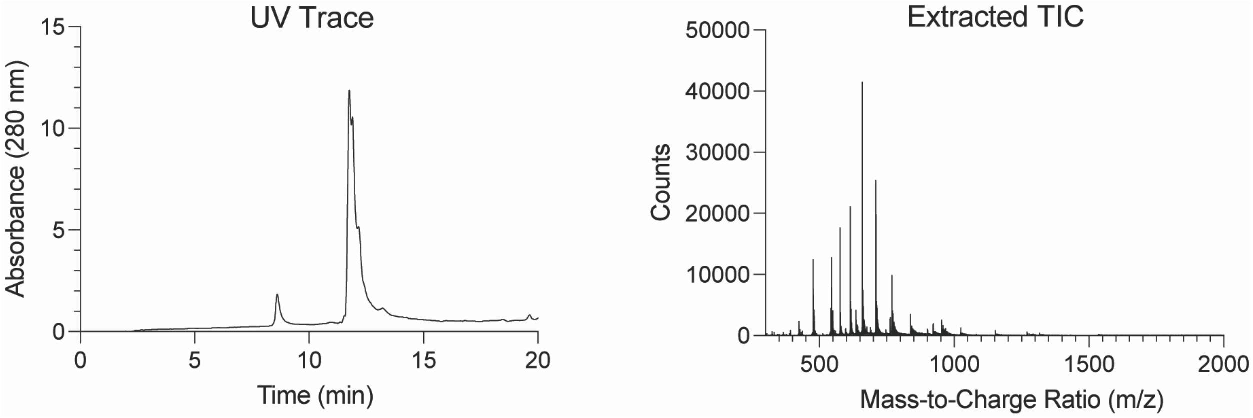

| | | |
|---|---|---|
| $L17ER_4$–PNA | Exp'd mass | 9213.85 |
| | Obs'd mass | 92.13.27 |

**Figure S6.** Representative deconvoluted Q–TOF mass spectra of $L17ER_4$–PNA.

#### R10–PNA

To conjugate the azide-containing R10 peptide to PNA-DBCO, a 1 mM stock solution of the R10–azide peptide and a 1 mM stock solution of the PNA-DBCO were prepared in water. The PNA and peptide were combined stoichiometrically in water. The reaction mixture was incubated for 48 h at 4 °C under agitation, but product formation plateaued. Accordingly, the reaction was heated to 50 °C for 6 h, and reaction progress was monitored by Q–TOF MS. The product was used directly without further purification.

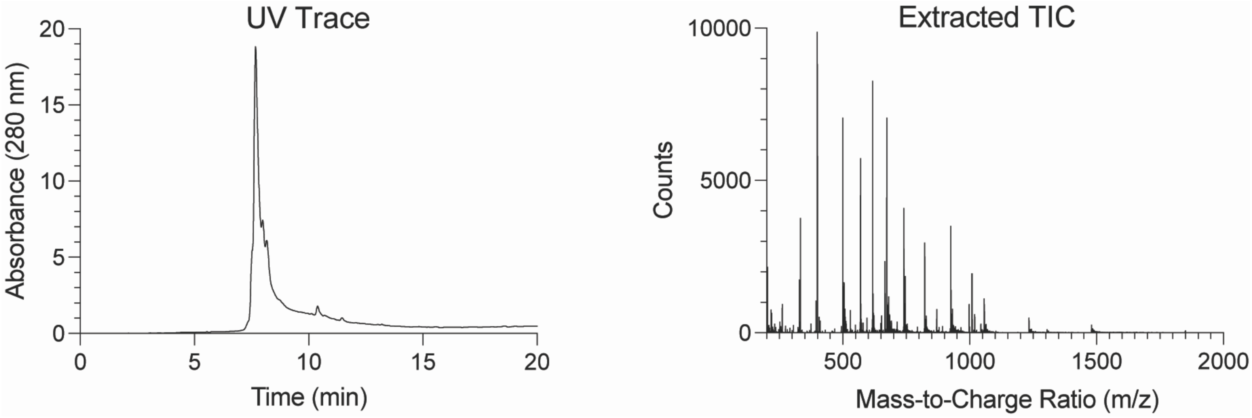

| | | |
|---|---|---|
| R10–PNA | Exp'd mass | 7397.77 |
| | Obs'd mass | 7398.56 |

**Figure S7.** Representative deconvoluted Q–TOF mass spectra of R10–PNA.

#### VI. Mammalian Cell Culture

HeLa654 cells (originally from the University of North Carolina Tissue culture Core facility) were cultured according to ATCC guidelines. HeLa654 cells were cultured in DMEM supplemented with FBS (10% v/v), penicillin (100 units/mL), and streptomycin (100 μg/mL). Cells were cultured in an incubator maintained at 37 °C and humidified to 5% v/v $CO_2$. Cells tested negative for mycoplasma.

#### VII. MTS Cytotoxicity Assay

The cytotoxicity of R10, L17E, and L17E$R_4$ was evaluated in HeLa654 cells using the CellTiter 96 AQueous One Solution Cell Proliferation Assay (product #G3580) from Promega (Madison, WI). This colorimetric assay leverages the metabolic activity of viable cells to convert a novel tetrazolium compound [3-(4,5-dimethylthiazol-2-yl)-5-(3-carboxymethoxyphenyl)-2-(4-sulfophenyl)-2*H*-tetrazolium, inner salt; MTS] into a colored, soluble formazan compound with absorbance in the visible light range. The absorbance of formazan produced is directly proportional to the living cell count.

Cells were seeded to be 90% confluent at the time of the experiment to most closely recapitulate treatment conditions in delivery experiments. Specifically, cells were seeded at 36,000 cells/well if performing the experiment 24 h later; cells were seeded at 18,000 cells/well if performing the experiment 48 h later; cells were seeded at 9,000 cells/well if performing the experiment 72 h later. Cells were seeded in a tissue culture treated flat black, clear bottom 96-well plate from Corning (product #3603). A stock solution of each peptide was prepared at 4 mM in water. A dilution series was prepared in either DMEM medium alone or full DMEM medium. The volume of vehicle in each treatment condition did not exceed 5% of the medium volume, and the working volume of each treatment condition was 100 μL. After treatment, cells were incubated for 16 h. Cells were then treated with 20 μL of the MTS reagent, and cells were incubated for 1.5 h at 37 ˚C and humidified to 5% v/v $CO_2$. Absorbance readings were collected at 490 nm on a Tecan Spark plate reader. Values represent data collected in biological duplicate with two technical replicates and are normalized to the average background absorbance signal from formazan in medium alone and from untreated cells. Data are plotted as the mean ± SE using Prism software from GraphPad (La Jolla, CA).

#### VII. Epifluorescence Microscopy of L17ER$_4$ for Morphology

Cells were seeded to be 90% confluent at the time of the experiment. Specifically, cells were seeded at 36,000 cells/well if performing the experiment 24 h later; cells were seeded at 18,000 cells/well if performing the experiment 48 h later; cells were seeded at 9,000 cells/well if performing the experiment 72 h later. In each case, cells were seeded into a sterile 18-well IbiTreat plate. Prior to treatment, cells were washed with DPBS without $Ca^{2+}/Mg^{2+}$ (2 × 100 µL), and serum-free or serum-containing medium was added to each well. The L17ER$_4$ peptide (1 mM stock in water) were prediluted to the appropriate treatment concentrations and subsequently dosed into each well according to the conditions below (Figure S8.) and incubated at 37 °C in a humidified incubator at 5% v/v $CO_2$. The volume of vehicle in each treatment condition did not exceed 5% of the medium volume, and the final volume of each well after the addition of peptide and PNA was equal to 50 µL. After incubation at 2-h and 24-h time points, cells were washed with DPBS without $Ca^{2+}/Mg^{2+}$ (2 × 100 µL) and Fluorobrite DMEM (100 µL) was used for imaging. Cells were protected from light at room temperature until imaged. Epifluorescent imaging was performed using an Evos M7000 Epifluorescent Microscope from ThermoFisher Scientific. The Transmitted light cube was used for excitation to assess cellular morphology and confluency. Images were collected using standardized laser intensity values. Images were analyzed using ImageJ software, adjusting for brightness and contrast, and processing was applied identically to all fluorescence images collected in a session.

#### Cytotoxicity Study with L17ER4

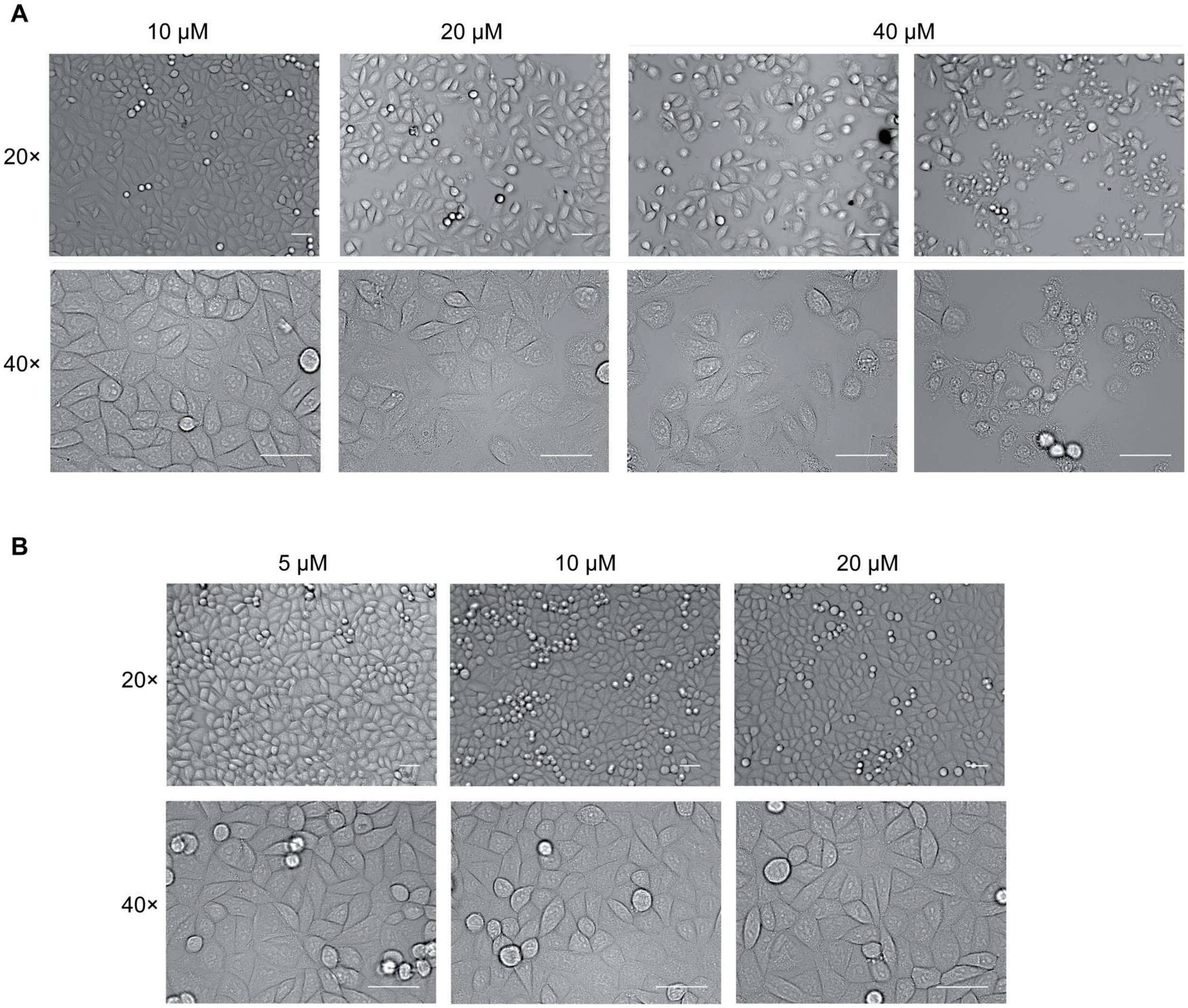

**Figure S8.** Transmitted light epifluorescence microscopy images of HeLa654 cells showing dose-dependent toxicity after treatment with L17ER4 after either (A) a 7-min treatment period in serum-free medium followed by a 2-h rest period at 37 °C in full medium, or (B) a 1-h treatment period in full medium followed by a 1-h rest period at 37 °C in full medium. Scale bars, 50 µm. All images were acquired with identical laser settings. Morphology and cell count was used to evaluate cell viability. The data in panel A demonstrate a cytotoxic effect from treatment at 40 µM, but robust viability at treatment concentrations at 20 µM and lower. The data in panel B demonstrate robust viability at all treatment conditions tested.

#### VIII. Epifluorescence Microscopy and Flow Cytometry for Fluorophore-Labelled Peptides

Cells were seeded to be 90% confluent at the time of the experiment. Specifically, cells were seeded at 36,000 cells/well if performing the experiment 24 h later; cells were seeded at 18,000 cells/well if performing the experiment 48 h later; cells were seeded at 9,000 cells/well if performing the experiment 72 h later. In each case, cells were seeded into a sterile 18-well IbiTreat plate. Prior to treatment, cells were washed with DPBS without $Ca^{2+}/Mg^{2+}$ (2 × 100 μL), and serum-free medium was added to each well. Peptide-fluorophore constructs were used directly in co-treatment experiments. All constructs were dissolved in water at 125 μM according to TAMRA fluorescence ($\varepsilon$ = 84,000 $M^{-1}$ $cm^{-1}$) as assessed with a DS-11 spectrophotometer. Conjugates were prediluted to the appropriate treatment concentrations in medium and subsequently dosed into each well according to the conditions below (Table S2) and incubated at 37 °C in a humidified incubator at 5% v/v $CO_2$. The volume of vehicle in each treatment condition did not exceed 5% of the medium volume, and the final volume of each well after the addition of fluorophore-labelled peptide was equal to 50 μL. After incubation, cells were washed with DPBS without $Ca^{2+}/Mg^{2+}$ (2 × 100 μL) lifted from the plate with 50 μL of 0.25% v/v trypsin–EDTA. Trypsin was quenched by the addition of 50 μL of full medium. Cells were strained into flow tubes and pelleted via centrifugation for 5 min at 1000 RPM at 4 °C. Cells were resuspended in 1 mL of ice-cold DPBS with $Ca^{2+}/Mg^{2+}$ supplemented with bovine serum albumin (0.1% w/v). Each sample was stained with SYTOX Blue Dead-Cell Indicator (1 μL of a 1.0 mM stock) for ≥5 min on ice protected from light. Cells were kept on ice and protected from light until the time of analysis. The fluorescence intensity of at least 10,000 live events was measured by flow cytometry with an Attune NxT flow cytometer (405 nm, 488 nm, 561 nm, and 640 nm lasers) from ThermoFisher Scientific. Control cells treated with SYTOX Blue Dead-Cell Indicator (1 μL of a 1.0 mM stock) were analyzed first to set gates and laser intensities. For the SYTOX Blue Dead-Cell Indicator, the 488-nm laser was used for excitation and the 530/30 filter was used to detect fluorescence. For TAMRA fluorescence, the 561-nm laser was used for excitation and the 585/16 filter was used to detect fluorescence. Events were collected using standardized laser intensity values. A template was established for laser intensity values, and all experiments were performed using these laser powers to enable comparison between data sets. Data were analyzed with FlowJo software, and the MFI of live, single cells is reported.

#### Table S2. Treatment Conditions for Fluorophore-Labelled Peptides

| Construct | Figure No. | Vehicle | Serum | Treatment Concentration (μM) | Treatment Time | Recovery Time |
|---|---|---|---|---|---|---|
| L17E–TAMRA | S9 and S10 | $H_2O$ | No | 5 | 5 min | 1 h |
| L17ER$_4$–TAMRA | S9 and S10 | $H_2O$ | No | 5 | 5 min | 1 h |
| TAMRA–L17E | S9 and S10 | $H_2O$ | No | 5 | 5 min | 1 h |
| TAMRA–L17ER$_4$ | S9 and S10 | $H_2O$ | No | 5 | 5 min | 1 h |

#### Representative Fluorophore-Labelled Peptide Uptake Data

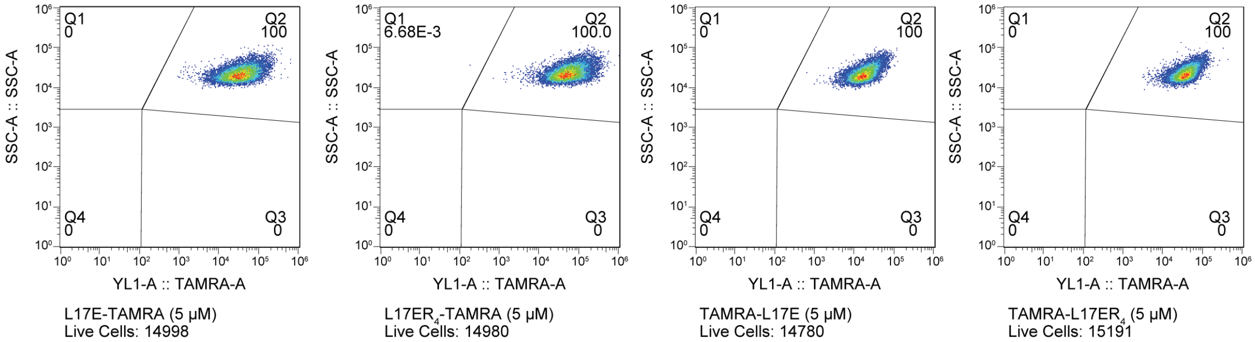

**Figure S9.** Flow cytometry of HeLa654 cells treated with 5 µM L17E or $L17ER_4$ labelled with TAMRA at either the N or C terminus. Representative flow plots are shown for each fluorophore-labelled peptide conjugate.

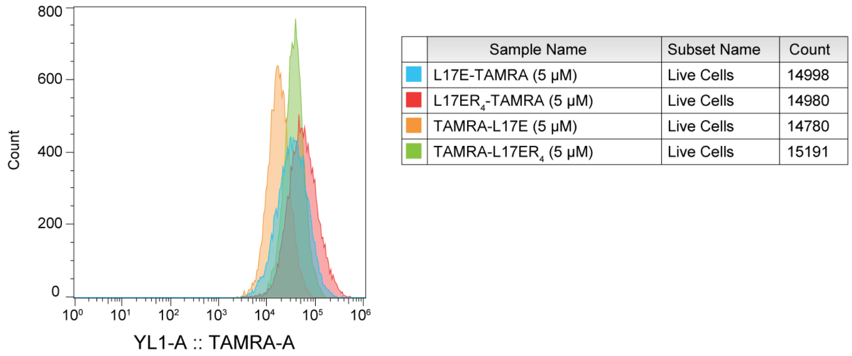

**Figure S10.** Histogram overlay of flow cytometry results from HeLa654 cells treated with 5 µM L17E or $L17ER_4$ labelled with TAMRA at either the N or C terminus. These data demonstrate that C-terminally modified peptides outperform N-terminally modified peptides.

#### IX. Epifluorescence Microscopy and Flow Cytometry for PNA Delivery

Cells were seeded to be 90% confluent at the time of the experiment. Specifically, cells were seeded at 36,000 cells/well if performing the experiment 24 h later; cells were seeded at 18,000 cells/well if performing the experiment 48 h later; cells were seeded at 9,000 cells/well if performing the experiment 72 h later. In each case, cells were seeded into a sterile 18-well IbiTreat plate. Prior to treatment, cells were washed with DPBS without $Ca^{2+}/Mg^{2+}$ (2 × 100 µL), and serum-free medium was added to each well. All constructs were used directly in co-treatment experiments. Constructs were prediluted in medium to the appropriate treatment concentrations and subsequently dosed into each well according to the conditions below (Table S3) and incubated at 37 °C in a humidified incubator at 5% v/v $CO_2$. The volume of vehicle in each treatment condition did not exceed 5% of the medium volume, and the final volume of each well after the addition of peptide and PNA was equal to 50 µL. After incubation, cells were washed with DPBS without $Ca^{2+}/Mg^{2+}$ (2 × 100 µL) and Fluorobrite DMEM (100 µL) was used for imaging. Cells were protected from light at room temperature until imaged. Epifluorescent imaging was performed using an Evos M7000 Epifluorescent Microscope from ThermoFisher Scientific. The GFP light cube ($\lambda_{ex}$ = 470/22 nm, $\lambda_{em}$ = 525/50 nm) was used for excitation. Images were collected using standardized laser intensity values. Images were analyzed using the open-source Fiji distribution of ImageJ, adjusting for brightness and contrast, and processing was identically applied to all fluorescence images collected in a session.

After imaging had concluded, cells were washed with DPBS without $Ca^{2+}/Mg^{2+}$ (2 × 100 µL) lifted from the plate with 50 µL of 0.25% v/v trypsin–EDTA. Trypsin was quenched by the addition of 50 µL of full medium. Cells were strained into flow tubes and pelleted via centrifugation for 5 min at 1000 RPM at 4 °C. Cells were resuspended in 1 mL of ice-cold DPBS with $Ca^{2+}/Mg^{2+}$ supplemented with bovine serum albumin (0.1% w/v). Each sample was stained with SYTOX Red Dead-Cell Indicator (1 µL of a 1.0 mM stock) for ≥5 min on ice protected from light. Cells were kept on ice and protected from light until the time of analysis. The fluorescence intensity of at least 10,000 live events was measured by flow cytometry with an Attune NxT Flow Cytometer (405 nm, 488 nm, 561 nm, and 640 nm lasers) from ThermoFisher Scientific. Control cells treated with SYTOX Red Dead-Cell Indicator (1 µL of a 1.0 mM stock), followed by cells treated with unmodified PNA, were analyzed first to set gates and laser intensities. For the SYTOX Red Dead-Cell Indicator, the 637-nm laser was used for excitation and the 670/14 filter was used to detect fluorescence. To detect GFP production, the 488-nm laser was used for excitation and the 530/30 filter was used to detect fluorescence. Events were collected using standardized laser intensity values. A template was established for laser intensity values, and all experiments were performed using these laser powers to enable comparison between data sets. Data were analyzed with the FlowJo software package (FlowJo), and the MFI of live, single cells is reported.

#### Table S3. Treatment Conditions for PNA Delivery

| Construct | Figure No. | Vehicle | Serum | Treatment Concentration (μM) | Treatment Time | Recovery Time |
|---|---|---|---|---|---|---|
| PNA654 + L17E | S14 and S15 | $H_2O$ | No | PNA654: 1, 5, 15, or 30 L17E: 40 | 5–10 min | 27 h |
| PNA654 + L17ER$_4$ | S16 and S17 | $H_2O$ | No | PNA654: 1, 5, 15, or 30 L17ER$_4$: 20 | 5–10 min | 27 h |
| PNA654 + R10 | S19 and S20 | $H_2O$ | No | PNA654: 1, 5, 15, or 30 R10: 40 | 5–10 min | 27 h |
| L17E–PNA654 | S21 and S22 | $H_2O$ | No | 5 or 10 | 5–10 min | 27 h |
| | | | Yes | 10 or 20 | 1 h | |
| L17ER$_4$–PNA654 | S23 and S24 | $H_2O$ | No | 5 or 10 | 5–10 min | 27 h |
| | | | Yes | 10 or 20 | 1 h | |
| R10–PNA654 | S25 and S26 | $H_2O$ | No | 5 or 10 | 5–10 min | 27 h |
| | | | Yes | 10 or 20 | 1 h | |

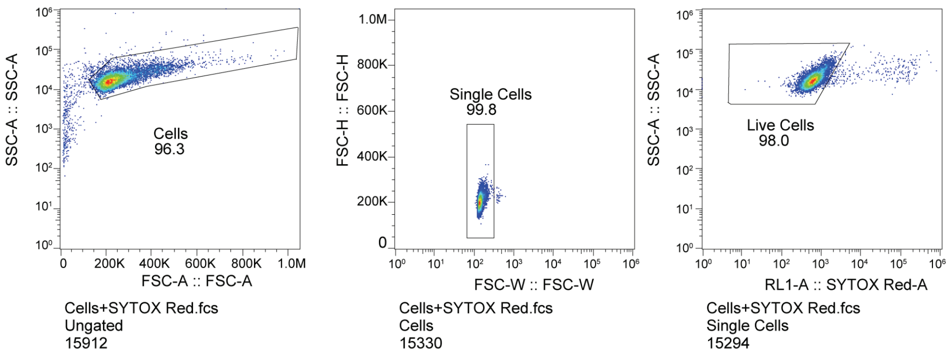

**Figure S11.** Representative flow cytometry gating strategy to identify live cells in a sample.

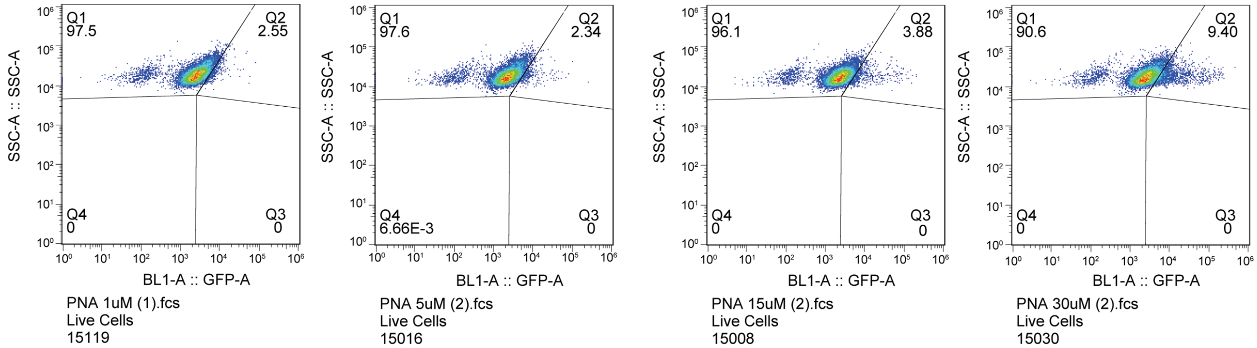

**Figure S12.** Flow cytometry of HeLa654 cells treated with 1, 5, 15, or 30 μM PNA654-SH. These data demonstrate that PNA alone has a low ability to enter the cytosol of cells to effect GFP production after corrective splicing.

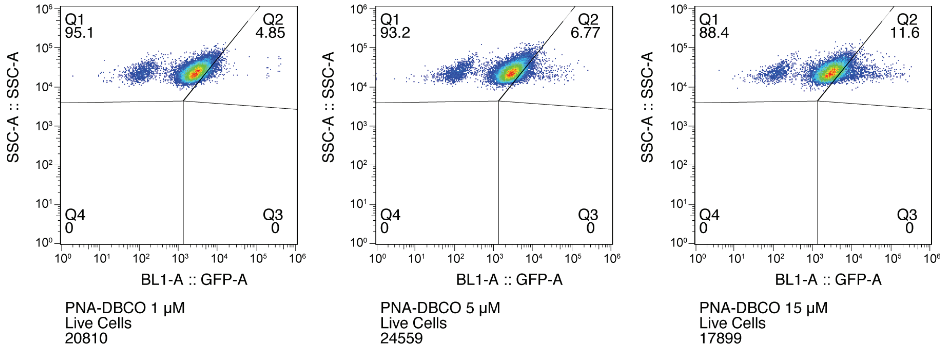

**Figure S13.** Flow cytometry of HeLa654 cells treated with 1, 5, or 15 μM PNA654-DBCO for 7 min in serum-free DMEM medium. These data demonstrate that PNA alone has a low ability to enter the cytosol of cells to effect GFP production after corrective splicing.

### Representative Co-treatment Data

#### L17E + PNA Co-treatment

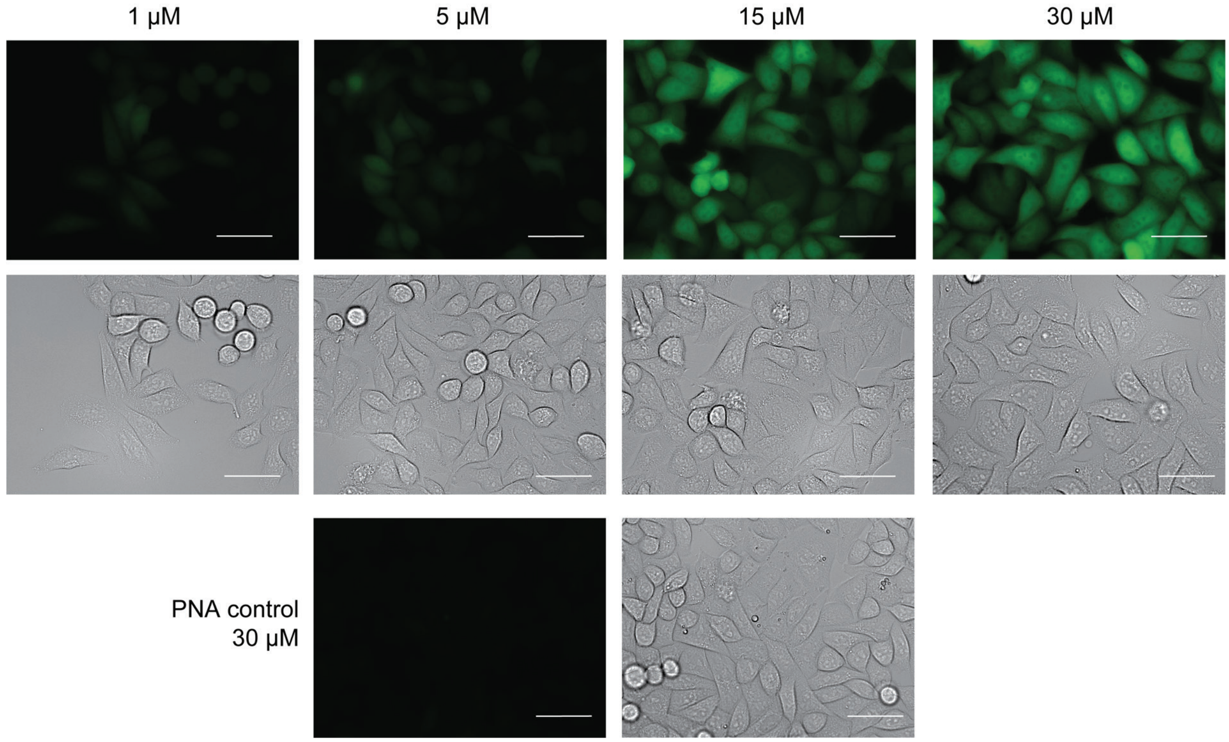

**Figure S14.** Fluorescence microscopy images of HeLa654 cells showing the dose-dependent GFP production after PNA co-treatment with L17E after a 7-min treatment period followed by a 27-h rest period at 37 °C. Scale bars, 50 µm. All images were acquired with identical laser settings.

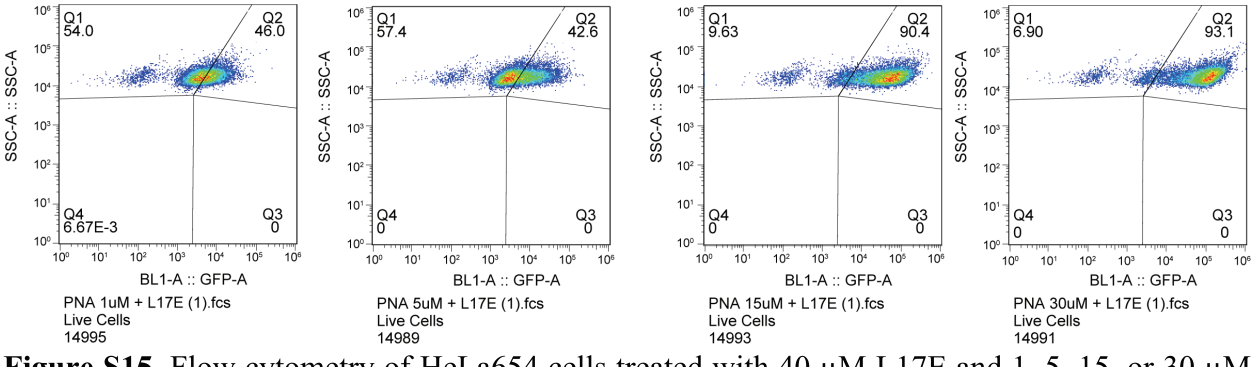

**Figure S15.** Flow cytometry of HeLa654 cells treated with 40 µM L17E and 1, 5, 15, or 30 µM PNA654-SH. These data demonstrate a robust dose–response of GFP production in cells that are treated with a higher concentration of PNA654-SH.

#### L17ER$_4$ + PNA Co-treatment

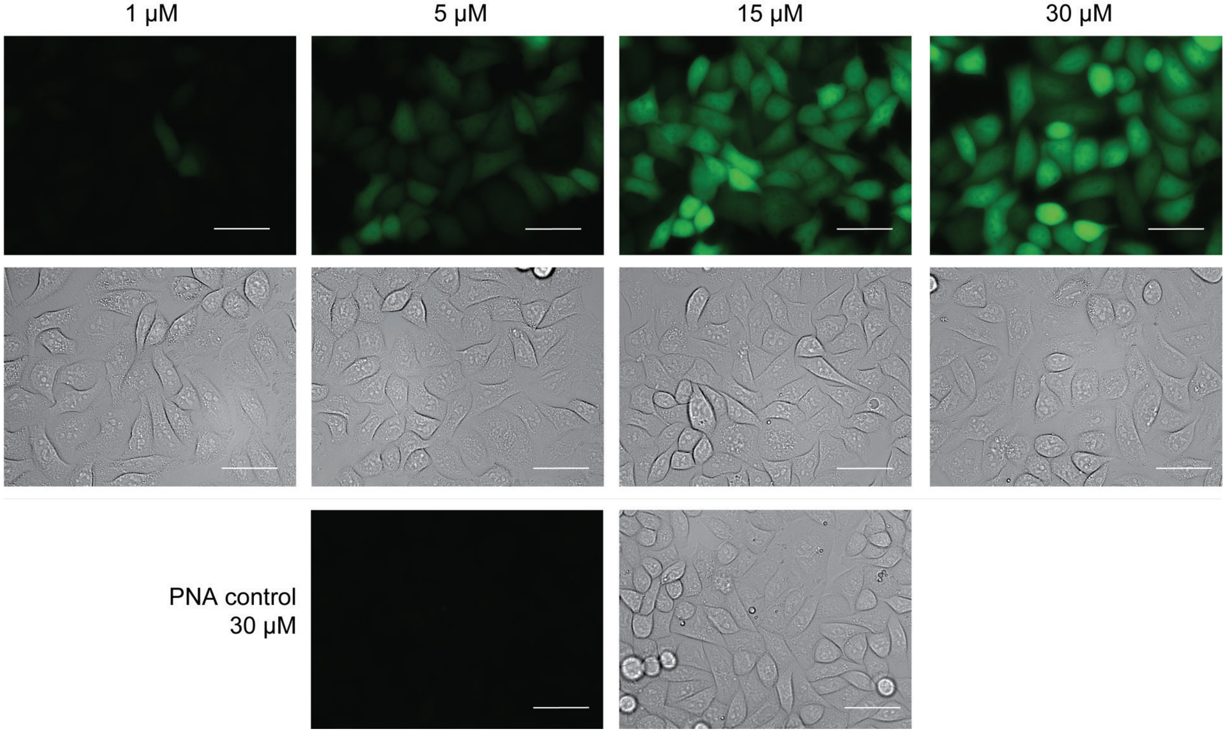

**Figure S16.** Fluorescence microscopy images of HeLa654 cells showing the dose-dependent GFP production after PNA co-treatment with L17ER$_4$ after a 7-min treatment period followed by a 27-h rest period at 37 °C. Scale bars, 50 μm. All images were acquired with identical laser settings.

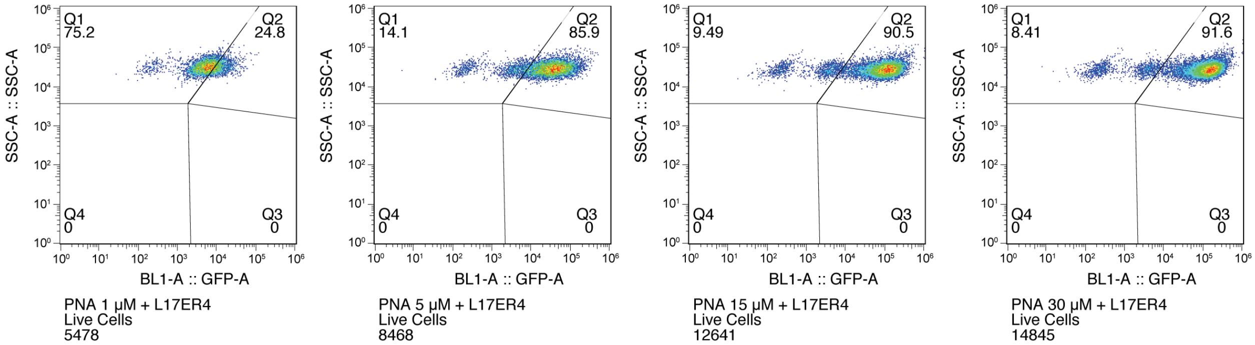

**Figure S17.** Flow cytometry of HeLa654 cells treated with 40 μM L17ER$_4$ and 1, 5, 15, or 30 μM PNA654-SH. These data demonstrate a robust dose–response of GFP production in cells that are treated with a higher concentration of PNA654-SH.

#### R10 + PNA Co-treatment

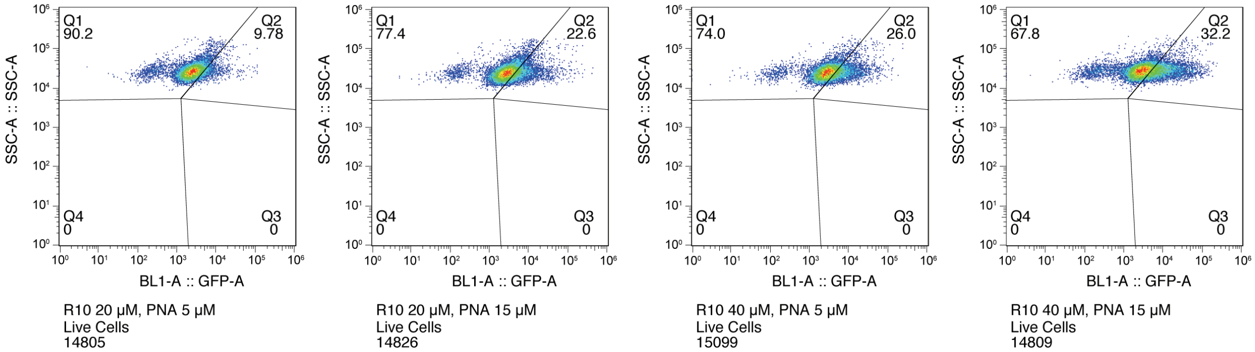

**Figure S18.** Flow cytometry of HeLa654 cells treated with 20 or 40 μM R10 and 5 or 15 μM PNA654-SH. The top row represents the universal gating strategy applied to all flow cytometry samples. These data demonstrate a weak degree of GFP production at either concentration of PNA654-SH, but that increasing the concentration of R10 increases GFP production. Because of this weak dose-dependent effect, 40 μM R10 was chosen for co-treatment experiments moving forward to provide a straightforward comparison to L17E.

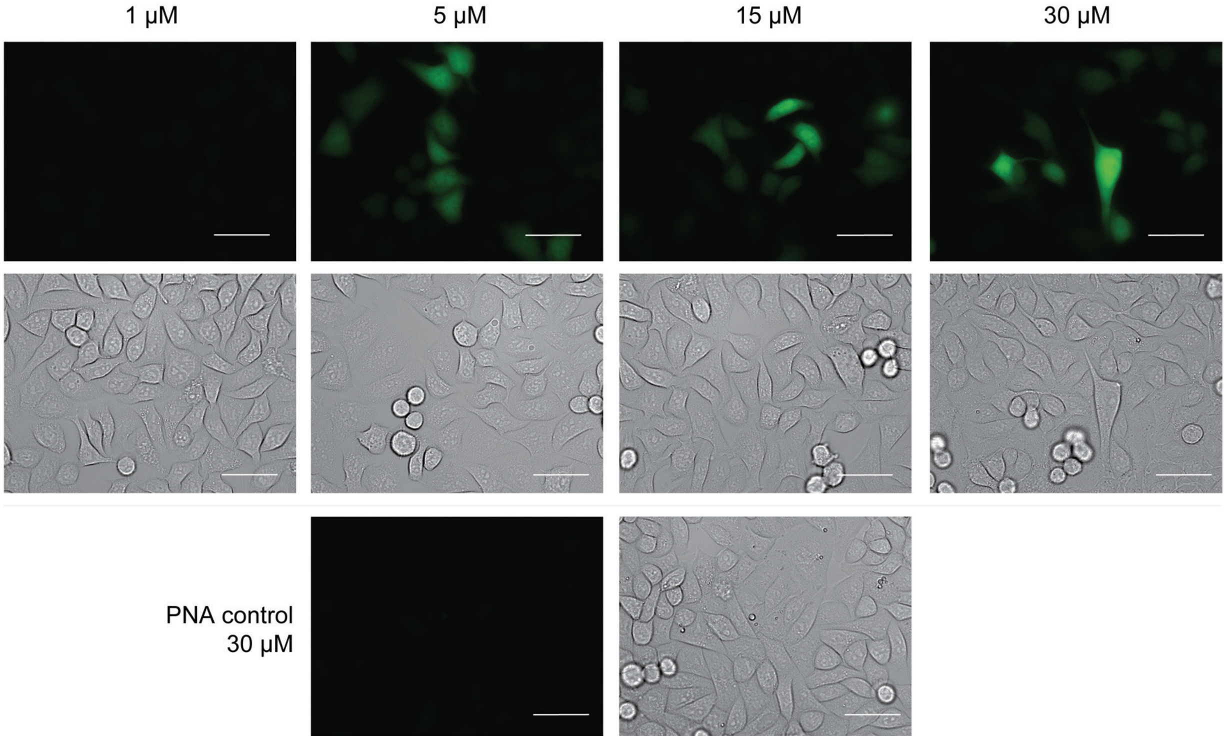

**Figure S19.** Fluorescence microscopy images of HeLa654 cells showing weak dose-dependent GFP production after PNA co-treatment with L17E after a 7-min treatment period followed by a 27-h rest period at 37 °C. Scale bars, 50 μm. All images were acquired with identical laser settings.

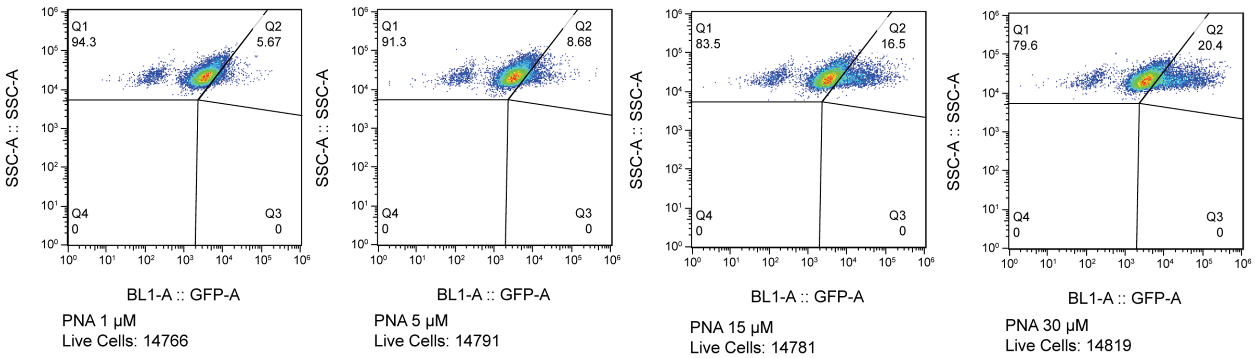

**Figure S20.** Flow cytometry of HeLa654 cells treated with 40 μM R10 and 1, 5, 15, or 30 μM PNA654-SH. These data demonstrate a weak dose response of GFP production in cells that are treated with a higher concentration of PNA654-SH.

#### Representative Covalent Irreversible Data

##### L17E–PNA

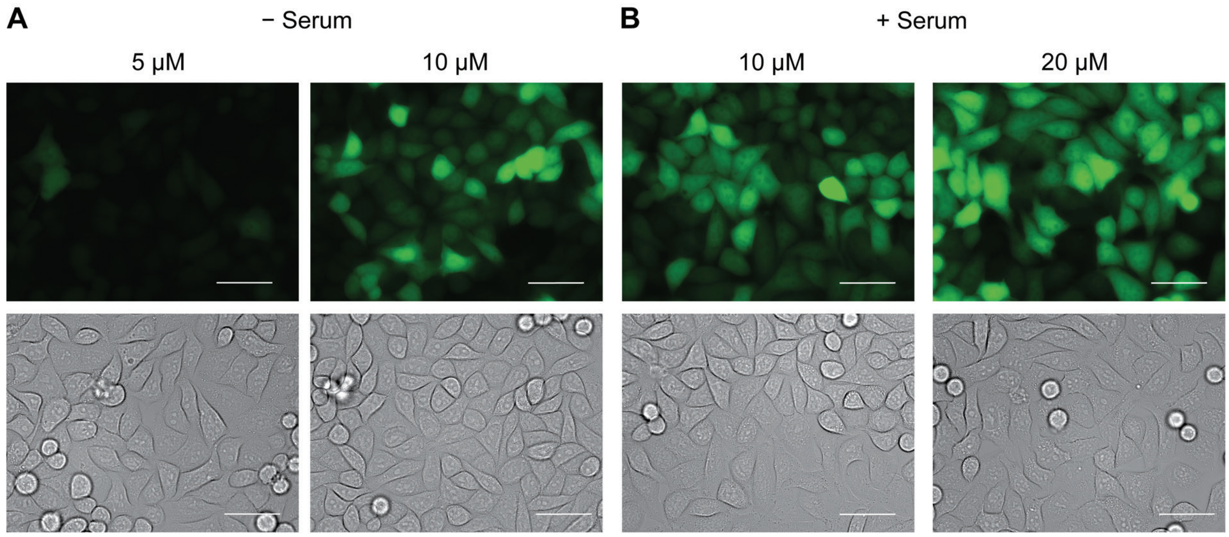

**Figure S21.** Fluorescence microscopy images of HeLa654 cells showing the dose-dependent GFP production after L17E-PNA treatment after (A) a 7-min treatment period in serum-free medium, or (B) a 1-h treatment in full medium followed by a 27-h rest period at 37 °C. Scale bars, 50 µm. All images were acquired with identical laser settings.

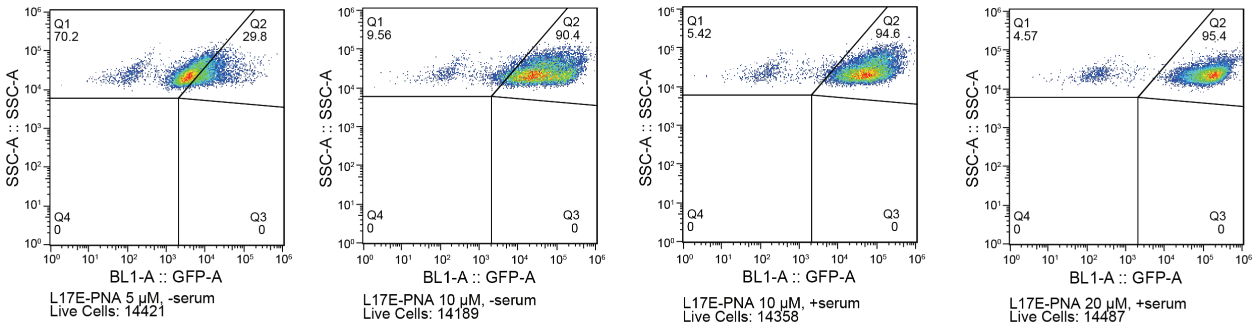

**Figure S22.** Flow cytometry of HeLa654 cells treated with L17E–PNA. These data demonstrate a robust dose–response of GFP production in cells that are treated with a higher concentration of L17E–PNA, both in the presence and absence of serum.

#### L17ER₄–PNA

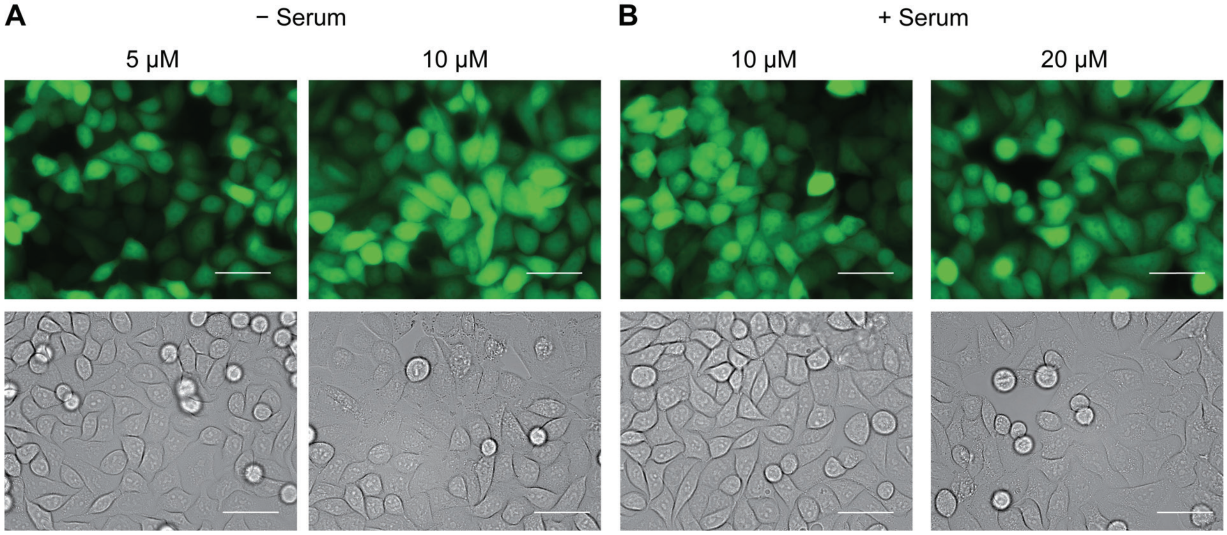

**Figure S23.** Fluorescence microscopy images of HeLa654 cells showing the dose-dependent GFP production after L17ER$_4$–PNA treatment after (A) a 7-min treatment period in serum-free medium, or (B) a 1-h treatment in full medium followed by a 27-h rest period at 37 °C. Scale bars, 50 µm. All images were acquired with identical laser settings.

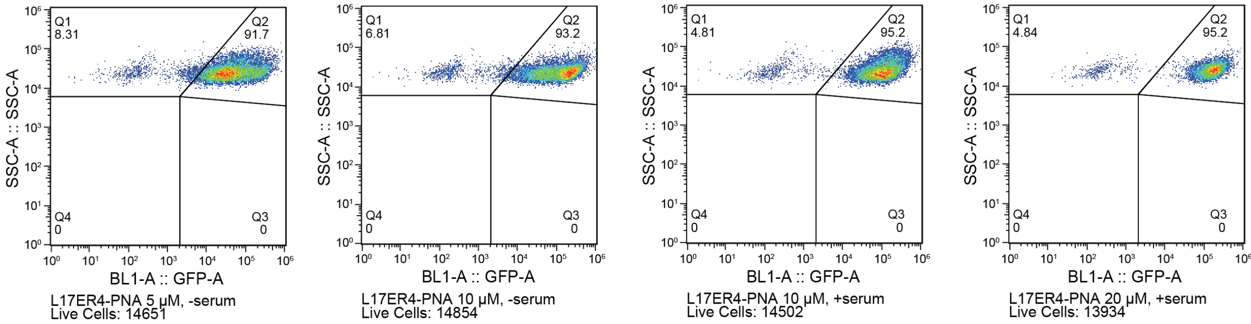

**Figure S24.** Flow cytometry of HeLa654 cells treated with L17ER$_4$–PNA. These data demonstrate a robust dose–response of GFP production in cells that are treated with a higher concentration of L17ER$_4$–PNA, both in the presence and absence of serum.

#### R10–PNA

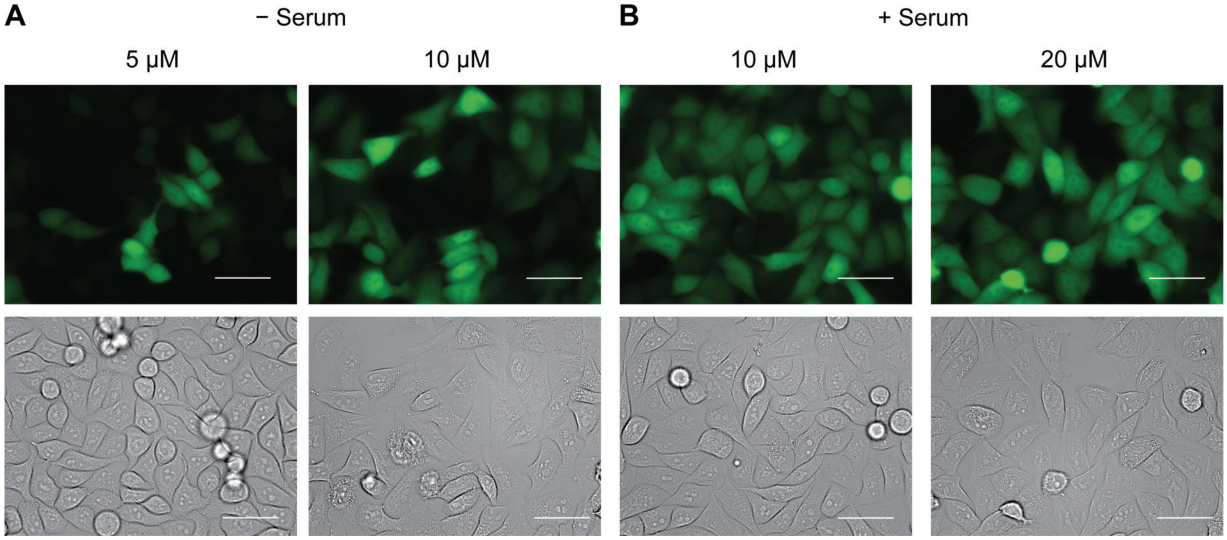

**Figure S25.** Fluorescence microscopy images of HeLa654 cells showing the dose-dependent GFP production after R10–PNA treatment after (A) a 7-min treatment period in serum-free medium, or (B) a 1-h treatment in full medium followed by a 27-h rest period at 37 °C. Scale bars, 50 μm. All images were acquired with identical laser settings.

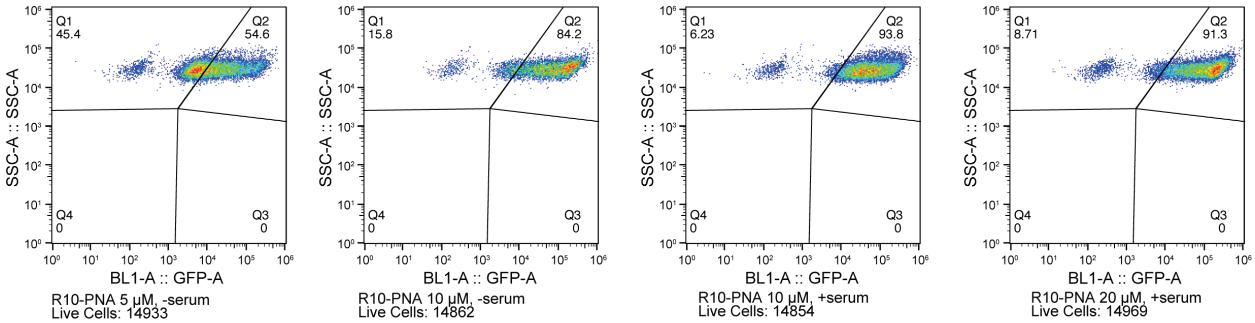

**Figure S26.** Flow cytometry of HeLa654 cells treated with R10–PNA. These data demonstrate a dose–response, both in the presence and absence of serum, though the cell populations demonstrate a higher spread of GFP production.

#### XI. Cell Synchronization Viability and PNA Delivery

Cells were seeded to be 90% confluent at the time of synchronization. Specifically, cells were seeded at 36,000 cells/well if performing the experiment 24 h later; cells were seeded at 18,000 cells/well if performing the experiment 48 h later; cells were seeded at 9,000 cells/well if performing the experiment 72 h later. In each case, cells were seeded into a sterile 18-well IbiTreat plate. The working volume of each well in the plate is 100 μL. To evaluate the impact of cell synchronization on cytosolic uptake, we evaluated synchronized and cycling cells side-by-side. At the time of confluency, half of the wells in the 18-well Ibidi dish were treated with RO-3306 (product #S7747) from Selleck Chemicals (Houston, TX), which is a reversible CDK1 inhibitor that arrests cells at the G2/M phase boundary. RO-3306 was reconstituted at a 20 mM concentration in DMSO. The stock solution was diluted to 2 mM in DMSO, and cells were treated at 9 μM for 20 h at 37 °C in a humidified incubator at 5% v/v $CO_2$. These treatment conditions are sufficient to synchronize cells without inducing cytotoxicity.

After 20 h, the viability of synchronized cells was evaluated compared to cycling cells in the same plate using the MTS assay as described previously, by the addition of 20 μL of the tetrazolium agent to 100 μL of culture medium. Cells were incubated for 1.5 h at 37 °C and humidified to 5% v/v $CO_2$. Absorbance readings were collected at 490 nm on a Spark plate reader. Values are normalized to the average background absorbance signal from formazan in medium alone and from cycling cells.

After 20 h, synchronized cells were released from arrest at the G2/M phase boundary by the removal of RO-3306 from the culture medium. When RO-3306 is withdrawn from the culture medium, cells begin cycling after ~40 min. Because cell division occurs over the course of ~1 h, synchronized and cycling cells were treated in serum-containing medium for 2 h to ensure that the period of cell division was captured. Synchronized and cycling cells were co-treated with L17E (40 μM) and PNA (5 or 15 μM), which were prediluted into full medium prior to addition to each sample well, for 2 h at 37 °C in a humidified incubator at 5% v/v $CO_2$. The volume of the vehicle in each treatment condition did not exceed 5% of the medium volume, and the final volume of each well after the addition of peptide and PNA was equal to 50 μL. After incubation, cells were washed with DPBS without $Ca^{2+}$/$Mg^{2+}$ (2 × 100 μL) lifted from the plate with 50 μL of 0.25% v/v trypsin–EDTA. Trypsin was quenched by the addition of 50 μL of full medium. Cells were strained into flow tubes and pelleted via centrifugation for 5 min at 1000 RPM at 4 °C. Cells were resuspended in 1 mL of ice-cold DPBS with $Ca^{2+}$/$Mg^{2+}$ supplemented with bovine serum albumin (0.1% w/v). Each sample was stained with SYTOX Red Dead-Cell Indicator (1 μL of a 1.0 mM stock) for ≥5 min on ice protected from light. Cells were kept on ice and protected from light until the time of analysis. The fluorescence intensity of at least 10,000 live events was measured by flow cytometry with an Attune NxT Flow Cytometer (405 nm, 488 nm, 561 nm, and 640 nm lasers) from ThermoFisher Scientific. Control cells treated with SYTOX Red Dead-Cell Indicator (1 μL of a 1.0 mM stock), followed by cells treated with unmodified PNA, were analyzed first to set gates and laser intensities. For the SYTOX Red Dead-Cell Indicator, the 637-nm laser was used for excitation and the 670/14 filter was used to detect fluorescence. For GFP production, the 488-nm laser was used for excitation and the 530/30 filter was used to detect fluorescence. Events were collected using standardized laser intensity values. A template was established for laser intensity values, and all experiments were performed using these laser powers to enable comparison between data sets. Data were analyzed with FlowJo software, and the MFI of live, single cells is reported.

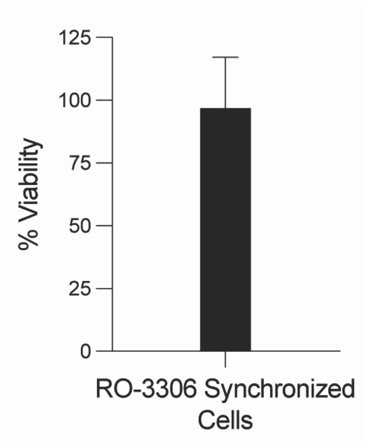

**Figure S27.** Graph of HeLa654 cell viability after treatment with RO-3306 (9 μM) for 20 h in a humidified incubator at 37 °C. Viability was assessed with a tetrazolium-based assay after 20 h. The values is the mean ± SE from one independent experiment, performed with two technical replicates.

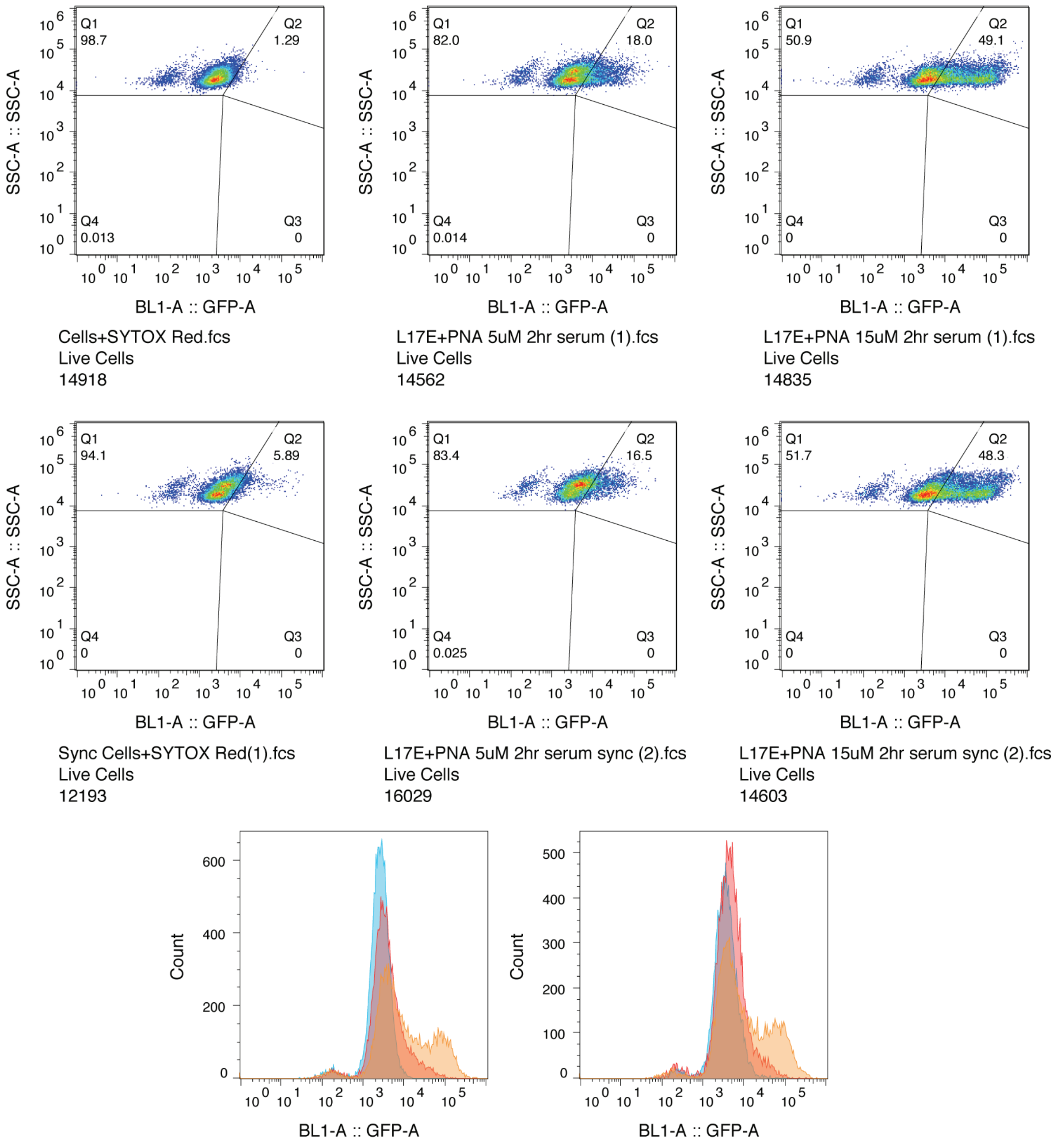

| Sample Name | Subset Name | Count |
|---|---|---|
| L17E+PNA 15uM 2hr serum (1).fcs | Live Cells | 14835 |
| L17E+PNA 5uM 2hr serum (1).fcs | Live Cells | 14562 |
| Cells+SYTOX Red.fcs | Live Cells | 14918 |

| Sample Name | Subset Name | Count |
|---|---|---|
| L17E+PNA 15uM 2hr serum sync (2).fcs | Live Cells | 14603 |
| L17E+PNA 5uM 2hr serum sync (2).fcs | Live Cells | 16029 |
| Sync Cells+SYTOX Red(1).fcs | Live Cells | 12193 |

**Figure S28.** Flow cytometry of HeLa654 cell GFP production after co-treatment with L17E (40 µM) and PNA-SH (5 or 15 µM) in cells that were cycling or cells that had been released from synchronization with RO-3306 (9 µM). The top row represents GFP production from cycling cells, and the middle row represents GFP production from synchronized cells. The histograms in the bottom row demonstrate that cycling cells have a higher degree of uptake and GFP production when compared to control than exhibited by synchronized cells.

#### XI. References

(1) Janson, N.; Krüger, T.; Karsten, L.; Boschanski, M.; Dierks, T.; Müller, K. M.; Sewald, N. Bifunctional reagents for formylglycine conjugation: Pitfalls and breakthroughs. *ChemBioChem* **2020**, *21* (24), 3580–3593. DOI: 10.1002/cbic.202000416.

(2) Debets, M. F.; van Berkel, S. S.; Schoffelen, S.; Rutjes, F. P. J. T.; van Hest, J. C. M.; van Delft, F. L. Aza-dibenzocyclooctynes for fast and efficient enzyme PEGylation via copper-free (3 + 2) cycloaddition. *Chem. Commun.* **2010**, *46* (1), 97–99. DOI: 10.1039/B917797C.
